# Lack of tau expression modifies epileptogenesis associated neuroplasticity in the ventral dentate gyrus in a mouse model of temporal lobe epilepsy

**DOI:** 10.64898/2026.09.23.753927

**Authors:** Madeleine C. Moseley, Bret N. Smith

**Author notes:** **Address Correspondence to:** Bret N. Smith, PhD,l Department of Biomedical Sciences Colorado State University, Fort Collins, CO 80523.

## Abstract

Deletion of the microtubule-associated protein, tau reduces seizure burden in rodent models of genetic epilepsies and suppresses evoked seizures. In the intrahippocampal kainate (IHK) model of acquired temporal lobe epilepsy (TLE), mice lacking tau expression develop spontaneous seizures, but at a lower rate than controls. Mechanisms by which tau deletion modifies the development of acquired TLE are not well understood, but a modification of the synaptic reorganization that accompanies epileptogenesis may contribute to modified seizure expression. Increased inhibitory synaptic input to dorsal dentate granule cells (DGCs) contralateral to the focal kainate injection results after development of TLE in tau^-/-^ mice, but inhibitory synaptic reorganization distal to the kainate lesion is undefined in this model. We therefore investigated how development of TLE modifies excitability and synaptic transmission in DGCs of the ventral dentate gyrus, distant from the kainate lesion. Our results show that development of TLE increases evoked action potential firing of ventral DGCs (vDGCs) in mice that develop TLE, demonstrating an effect of TLE development on excitability of DGCs distant from the focal injury. After development of TLE, DGCs from tau^-/-^ mice received increased inhibitory synaptic input bilaterally, suggesting that tau expression suppresses inhibitory synaptic reorganization in DGCs. Optogenetic stimulation of interneurons in tau^-/-^ mice revealed increased inhibitory synaptic input to vDGCs and an increase in inhibitory convergence onto vDGCs once epileptogenesis occurred. Therefore, tau deletion modifies spontaneous seizure expression after epileptogenesis in mice, potentially involving increased synaptic inhibition of DGCs throughout the dentate gyrus.

## Introduction

Studies in which the microtubule associated protein tau is genetically deleted or suppressed in models of genetic epilepsies (i.e., channelopathies) and chemoconvulsant-evoked seizures demonstrate that lack of tau expression is protective against seizure expression (DeVos *et al*., 2013; Holth *et al*., 2013; Gheyara *et al*., 2014; Li *et al*., 2014; Putra *et al*., 2020; Shao *et al*., 2022), but studies examining how deletion of tau modifies epileptogenesis in acquired TLE models are limited. In the intrahippocampal kainate (IHK) model of acquired TLE, lack of tau expression is not effective in preventing epileptogenesis or the development of spontaneous seizures in acquired TLE but influences seizure expression and network activity during the ictal phase of seizures (Moseley *et al*., 2025).

In the IHK model of acquired TLE, focal injection of kainic acid into the dorsal hippocampus typically results in development of status epilepticus (SE) and after a delay of 2-3 weeks, mice develop unprovoked spontaneous seizures. The development of spontaneous recurrent seizures in the IHK and other rodent models of TLE is often associated with pathological hallmarks that recapitulate characteristics seen in human patients with TLE such as loss of subsets of inhibitory interneurons, dentate granule cell (DGC) axon sprouting, synaptic reorganization of DGC connections, and DGC dispersion (deLanerolle *et al*., 1989; Sutula *et al*., 1989; Leite *et al*., 1996; Bouilleret *et al*., 1999; Sundstrom *et al*., 2001; deLanerolle *et al*., 2003; Sutula & Dudek, 2007; Marx *et al*., 2013; Sieu *et al*., 2017; Zeidler *et al*., 2018; Kang *et al*., 2021; Lee *et al*., 2024; Moseley *et al*., 2025).

Unlike TLE induced after intraperitoneal chemoconvulsant administration, acute neuropathology in the unilateral IHK model is largely limited to the ipsilateral hippocampus with distant sites remaining relatively intact. Synaptic reorganization that is limited to the dorsal hippocamps however, may be ineffective in regulating or inducing seizures, and seizures in other acquired TLE models and patients with TLE often develop in the ventral hippocampus (J. Engel *et al*., 1990; Buckmaster *et al*., 2022). The focal IHK model allows for the investigation of cellular and synaptic changes associated with epileptogenesis in brain regions remote from the injury, like the ventral hippocampus (Bouilleret *et al*., 1999; Riban *et al*., 2002; Gouder *et al*., 2003; Groticke *et al*., 2008; Zeidler *et al*., 2018; Kang *et al*., 2021). While mice lacking tau expression (i.e., tau^-/-^ mice) developed TLE at a lower rate than wildtype controls, with reduced severity of spontaneous seizures, development of TLE in tau^-/-^ mice was additionally associated with increased inhibitory synaptic drive to DGCs contralateral to the kainate injection in the dorsal hippocampus (Moseley et al., 2025). This suggests discrete inhibitory synaptic reorganization may be a potential mechanism by which tau deletion suppresses seizure expression once epilepsy develops. Therefore, we sought to examine how a focal insult (kainate injection) in the dorsal hippocampus alters epileptogenesis-associated neuroplasticity in the ventral hippocampus, an area remote from the insult, and if lack of tau expression modifies neuroplastic changes associated with the development of TLE.

We assessed neuronal excitability and synaptic transmission in ventral DGCs, not only because of their involvement in epileptogenesis and tauopathy-associated pathology, but also to investigate if epileptogenesis associated neuroplasticity is evident at sites distal from the focal lesion in the dorsal hippocampus in mice lacking tau expression and in wildtype C57Bl/6J controls. We tested the hypothesis that tau deletion contributes to the synaptic reorganization of inhibitory circuits during epileptogenesis in acquired TLE.

## Material and Methods

### Animals

Transgenic B6.Cg-Mapttm1(EGFP)KltTg(MAPT)8cPdav/J mice were produced in house from breeders obtained from The Jackson Laboratory (JAX; Bar Harbor, ME; stock #005491). All breeding mice were homozygous for the deletion of the murine tau gene. One mouse in each breeding pair was hemizygous for a transgene expressing all six isoforms of non-mutant, human tau protein. The offspring are therefore either full tau knockout (tau^-/-^) or express only human tau (htau), and genotypes and protein expression phenotypes were confirmed in our recent reports (Cloyd et al., 2021, Moseley et al., 2025). Here, we used tau^-/-^ mice and isogenic C57BL/6J wildtype control mice that express murine tau protein.

Age matched male C57BL/6J control mice were bred in house from breeders obtained from JAX (#000664). All mice were housed under a 12 hr light / 12 hr dark cycle in an Association for Assessment and Accreditation of Laboratory Animal Care (AAALAC) approved facility. Food and water were available ad libitum. The Colorado State University Laboratory Animal and Use Committee approved all procedures. Principles outlined in the Animal Research: Reporting of in Vivo Experiments (ARRIVE) guidelines and the Basel declaration, including the 3R (Replacement, Reduction, Refinement) concept, were considered when planning the experiments.

### Intrahippocampal kainate model of TLE

All surgical procedures were performed under isoflurane general anesthesia with 0.05% bupivacaine local anesthesia. Kainic acid (120 nL, 20 mM in 0.9% saline, Tocris Bioscience; Minneapolis, MN) or saline (sham; 120 nL, 0.9% saline) was injected into the left dorsal hippocampus in male and female tau^-/-^ and wildtype mice between 8 and 10 weeks of age (Krook-Magnuson *et al*., 2013), as described previously (Moseley *et al*., 2025). After injection, mice were monitored for status epilepticus (SE) and diazepam (7.5 mg/kg) was administered intraperitoneally to terminate SE 4 hours after injection.

Development of TLE with the expression of spontaneous seizures in mice was determined by a continuous 72 hr video recording session at 5-6 weeks after the IHK or vehicle injection. Mice lacking tau expression develop TLE but at a lower rate than non-transgenic controls in this model (Moseley *et al*., 2025). All IHK-treated mice used in this study exhibited at least one spontaneous seizure during the observed video-recording period and were considered to have developed TLE. We included both males and females from both genotypes and treatments (Vehicle: WT: Males N=7, Females N=11; tau^-/-^: Males N=9, Females N=8. IHK: WT: Males N=6, Females N=8; tau^-/-^: Males N=10, Females N=6).

### Adeno-associated viral injections in the ventral hippocampus

In the same surgery as saline or IHK injections, 500 nL of rAAV-mDlx-ChR2-mCherry-Fishell-3 (Addgene; #83898-AAV9) was injected bilaterally into the ventral hippocampus (-3.5 mm posterior; ±2.8 mm medial/lateral, -3.8 mm ventral). The injection rate was 10 nL/min, and the needle (Hamilton syringe) was left in place 5 min before and after injection.

### Whole-cell patch clamp electrophysiological recordings

Horizontal slices containing the ventral hippocampus or coronal slices containing the dorsal hippocampus (300 µm) from vehicle and IHK-treated tau^-/-^ and wildtype mice 6-8 weeks after IHK or saline injection were prepared in cold, oxygenated sucrose artificial cerebrospinal fluid (ACSF; in mM: 85 NaCl, 75 Sucrose, 2.5 KCl, 25 D-glucose, 1.25 NaH_2_PO_4_, 4 MgCl_2_, 0.5 CaCl_2_, 24 NaHCO_3_). Slices were incubated for one hour at 34°C then transferred to a recording chamber containing ACSF (in mM): 124 NaCl, 26 NaHCO_3_, 11 glucose, 3 KCl, 2 CaCl_2_, 1.3 MgCl_2_, and 1.4 NaH_2_PO_4_; 32-34°C for recordings.

Whole-cell patch-clamp recordings were obtained from ventral DGCs (vDGCs) and dorsal DGCs (dDGCs) identified by location within the granule cell layer and morphological characteristics. Electrophysiological recordings were performed as previously described (Moseley *et al*., 2025) using a Multiclamp 700B amplifier (Molecular Devices, San Jose, CA) and acquired using the pClamp software suite (Molecular Devices). Recordings were discarded if series resistance changed by more than 20% during the recording or reached 25 MΩ. Recording pipettes were pulled from borosilicate glass (open tip resistance 3-5 MΩ; King Precision Glass Co., Claremont, CA). Current clamp recordings were obtained using a pipette recording solution which contained (in mM) 130 K-gluconate, 10 HEPES, 1 NaCl, 2 MgATP, 1 MgCl_2_, 1 CaCl_2_, and 5 EGTA; pH adjusted to 7.2 with KOH with an osmolarity of 290-295 mOsm. Voltage clamp recordings were obtained using a pipette recording solution which contained (in mM) 140 Cs-gluconate, 10 HEPES, 1 NaCl, 2 MgATP, 1 MgCl_2_, 1 CaCl_2_, and 5 EGTA; pH adjusted to 7.2 with CsOH with an osmolarity of 290-295 mOsm. Liquid junction potential was calculated using JPCalc (pClamp, Molecular Devices) to be +15.56 mV for K-gluconate and +16.57 mV for Cs-gluconate internals.

Intrinsic properties of vDGCs and dDGCs were recorded in current clamp configuration and membrane properties assessed using 1 s hyperpolarizing and depolarizing current steps of 20 pA increments from -200 pA to 200 pA from a membrane potential of -70 mV. Resting membrane potential was recorded in I=0 configuration and averaged over 60 seconds. Input resistance was measured using currents steps from -20 to 20 pA. Rheobase was defined as the first depolarizing current step that elicited action potentials. Current clamp recordings were analyzed using Clampfit 11.2 (Molecular Devices). Synaptic properties of vDGCs were measured by recording spontaneous excitatory post-synaptic currents (sEPSCs) at a holding potential of -70 mV and spontaneous inhibitory post-synaptic currents (sIPSCs) at a holding potential of 0 mV in voltage-clamp configuration. Action potential independent, miniature excitatory post-synaptic currents (mEPSCs) and miniature inhibitory post-synaptic currents (mIPSCs) were recorded as spontaneous events in the presence of 1 µM tetrodotoxin (TTX, Alomone Labs). Voltage clamp recordings were analyzed using Mini Analysis (version 6.0.7, Synaptosoft Inc.). Synaptic current amplitude was required to be greater than three times root mean square noise level to be included in s/mEPSCs and s/mIPSC frequency and amplitude analyses.

To examine local inhibitory interneuron mediated synaptic connections with vDGCs, we used mice expressing pAAV-mDlx-Ch2-mCherry to identify and optogenetically stimulate inhibitory interneurons while recording vDGCs in the presence of 1 mM kynurenic acid (Sigma-Aldrich) to block ionotropic glutamate receptor-mediated synapses. While recording vDGCs, mCherry-Ch2 expressing interneurons were activated using blue light centered on the soma of an identified interneuron through the 40x objective. To limit the simulation of more than one mCherry expressing interneuron and the dendrites or axons of other interneurons, the light field stop was closed to ensure light activation of a diameter of ∼30 µm. Evoked inhibitory post-synaptic currents (eIPSCs) were measured at 0 mV. To evaluate a successful connection between an interneuron and vDGCs, 5 light stimuli (10 ms duration, 1 sec inter-stimulus interval) were delivered to an interneuron while recording a vDGC (Supplemental Figure 1). A successful connection was defined as at least 3 eIPSCs per 5 stimulations as previously described (Hunt *et al*., 2010). Stimulations of mCherry expressing interneurons within 60 µm of the recorded vDGC (i.e., twice the stimulus radius) were discarded to avoid analysis of responses to direct stimulation of the recorded vDGC. Connection map schematics were created using the recorded micron location of vDGCs and interneurons using Matlab 2026a. The average peak eIPSC amplitude was calculated for successful connections across sequential stimulations using Clampfit 11.2 (Molecular Devices). Charge transfer (pA x ms) and decay time (ms) of eIPSCs were analyzed using Matlab R2026a. To determine short-term plasticity of GABAergic synapses from connecting interneurons and vDGCs, interneurons were stimulated with higher frequency light stimuli trains (10 light stimulations, 2 ms duration, 10 ms interstimulus interval). Percent change in eIPSC amplitude was measured using Clampfit where evoked IPSCs with an amplitude greater than 30 pA were included as evoked responses. In addition, efficacy of optogenetic activation of mCherry expressing interneurons were measured in current clamp to verify the equivalence of stimulus response across genotypes (Supplemental Fig. 2). Action potentials were evoked in mCherry-expressing interneurons from -60 mV by 5 light stimuli (10 ms duration, 1 second interval).

Excitatory synaptic input to mCherry expressing interneurons was measured in voltage clamp. Spontaneous excitatory post synaptic currents (sEPSCs) were recorded at a holding potential of -70 mV. Recordings were analyzed using Mini Analysis (version 6.0.7, Synaptosoft Inc.). Synaptic current amplitude was required to be greater than three times root mean square noise level to be included in sEPSCs frequency and amplitude analyses.

### Timm’s Stain of Mossy Fiber Sprouting

Ventral hippocampal slices used for recordings were incubated in 0.37% sodium sulfide solution for 20 minutes followed by 4% paraformaldehyde in 0.15 M phosphate buffer overnight. Slices were washed 3 times for 10 minutes in 0.01 M phosphate buffered saline (PBS) and cryoprotected overnight in 30% sucrose solution in PBS. Slices were sectioned at 20 µM, rinsed in 0.15 M phosphate buffer and mounted on charged slides (Superfrost Plus; Fisher Scientific) and dried for 48 hours. Sections were developed as previously described (Hunt *et al*., 2010; Hunt *et al*., 2012; Butler *et al*., 2015; Kang *et al*., 2022). Timm’s scoring was performed by two investigators blinded to treatment and genotype.

### Statistical Analysis

Statistical measures were performed with Prism (GraphPad, San Diego, CA). Electrophysiological data were analyzed using one-way ANOVA when comparing treatments within genotypes (Figs. 6-8) and two-way ANOVA when comparing across genotypes and treatments (Figs. 1-5) with Tukey post-hoc test for multiple comparisons when appropriate. Data are presented as mean ± SEM and statistical significance was set to p < 0.05 for all tests. Statistical tests are shown in the Statistics Table (Table 1).

**Figure 1.**
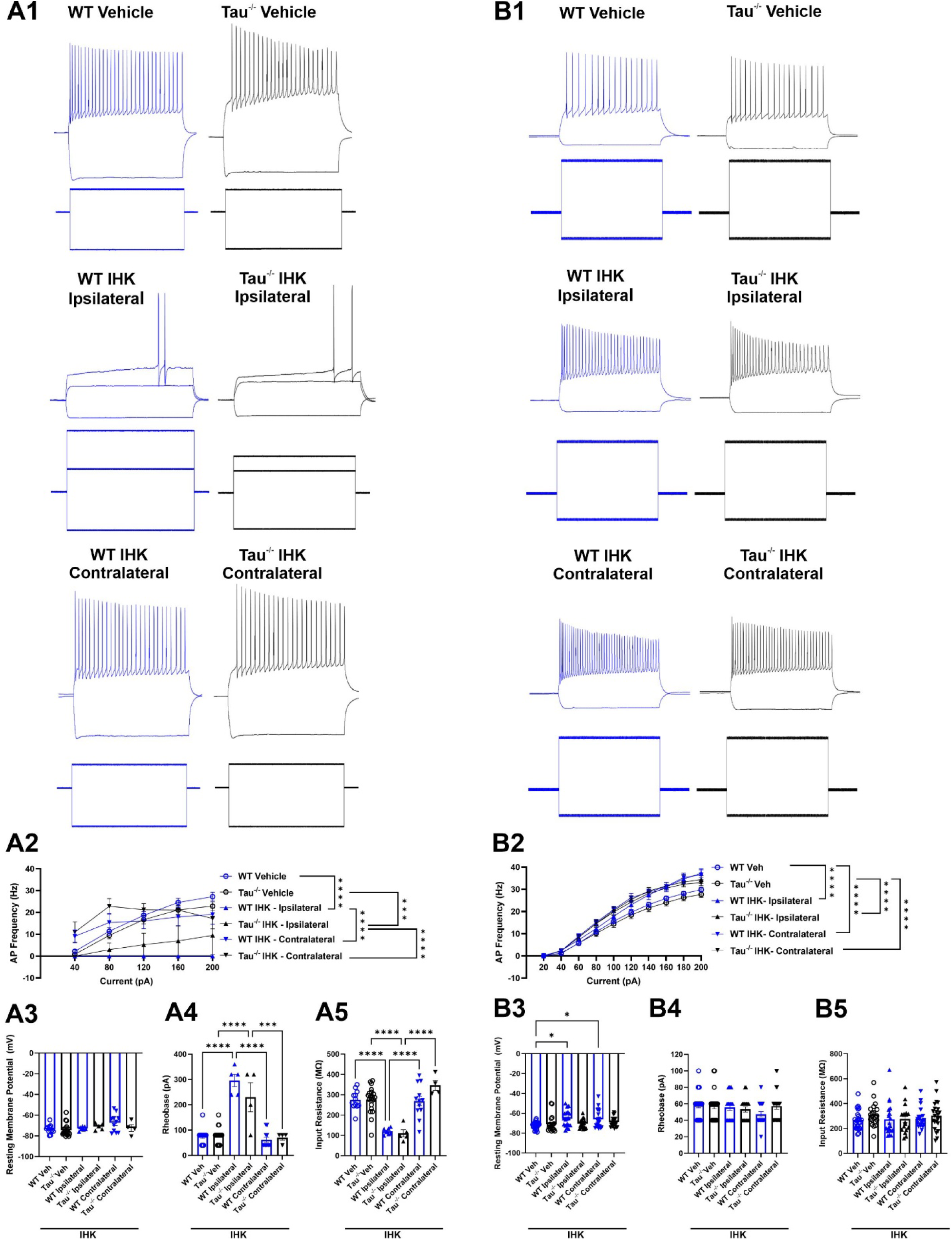
Intrinsic excitability is reduced in dorsal DGCs near IHK injection and increased in ventral DGCsin mice that develop TLE. (A) Action potential firing of dorsal dentate granule cells (dDGCs) in response to current injection steps (A1) dDGCs recorded from wildtype (WT) and tau^-/-^ vehicle controls demonstrate voltage responses at -200 pA and 120 pA. dDGCs ipsilateral to intrahippocampal kainate (IHK) injection in WT and tau^-/-^ mice. WT IHK-ipsilateral trace demonstrates voltage responses at -200 pA, 120 pA, and 240 pA. Rheobase of this cell was at 240 pA. Tau^-/-^ IHK-ipsilateral demonstrates voltage responses at -200 pA, 120 pA, 200 pA. Rheobase of this cell was at 200 pA. dDGCs contralateral to IHK injection in WT and tau^-/-^ mice demonstrate voltage responses at -200 pA and 120 pA. (A2) Input-out curve of action potential frequency of dDGCs from vehicle and IHK-treated wildtype and tau^-/-^ mice (***p<0.001, ****p<0.0001, Two-way ANOVA). (A3) Resting membrane potential of dDGCs from vehicle and IHK-treated wildtype and tau^-/-^ mice (p>0.05, Two-way ANOVA). (A4) Rheobase (i.e., minimum current that elicits action potentials) of dDGCs from vehicle and IHK-treated wildtype and tau^-/-^ mice (***p<0.001, ****p<0.0001, Two-way ANOVA). (A5) Input resistance of dDGCs from vehicle and IHK-treated wildtype and tau^-/-^ mice (***p<0.0001; Two-way ANOVA). WT Vehicle: N=4, n=19; Tau^-/-^ Vehicle: N=4, n=21; WT IHK: N=2, ipsilateral: n=5, contralateral: n=11; Tau^-/-^ IHK N=3, ipsilateral: n=5, contralateral: n=4). (B) Action potential firing of vDGCs in response to current injection steps. (B1) Representative traces of evoked action potentials in response to current injection steps from vehicle and IHK-treated WT and tau^-/-^ mice. (B2) Input-output curves of action potential frequencies for vDGCs from vehicle and IHK-treated WT and tau^-/-^ mice (*, different from vehicle-treated wildtype controls, +, different from vehicle-treated tau^-/-^ mice; p<0.05, Two-way ANOVA). (B3) Resting membrane potential of vDGCs from vehicle and IHK-treated WT and tau^-/-^ mice (p>0.05, Two-way ANOVA). (B4) Rheobase (i.e., minimum current that elicits action potentials) of vDGCs from vehicle and IHK-treated WT and tau^-/-^ mice (p>0.05, Two-way ANOVA). (B5) Input resistance of vDGCs from vehicle and IHK-treated WT and tau^-/-^ mice. (p>0.05, Two-way ANOVA). WT Vehicle: N=7, n=25; Tau^-/-^ Vehicle: N=6, n=20; WT IHK: N=10, ipsilateral: n=19, contralateral: n=20; Tau^-/-^ IHK N=11, ipsilateral: n=18, contralateral: n=22).

**Table 1.**
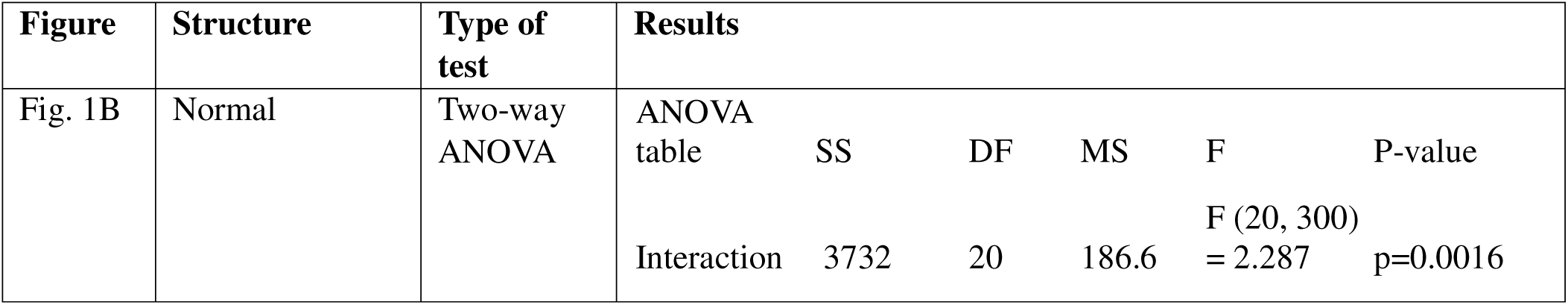
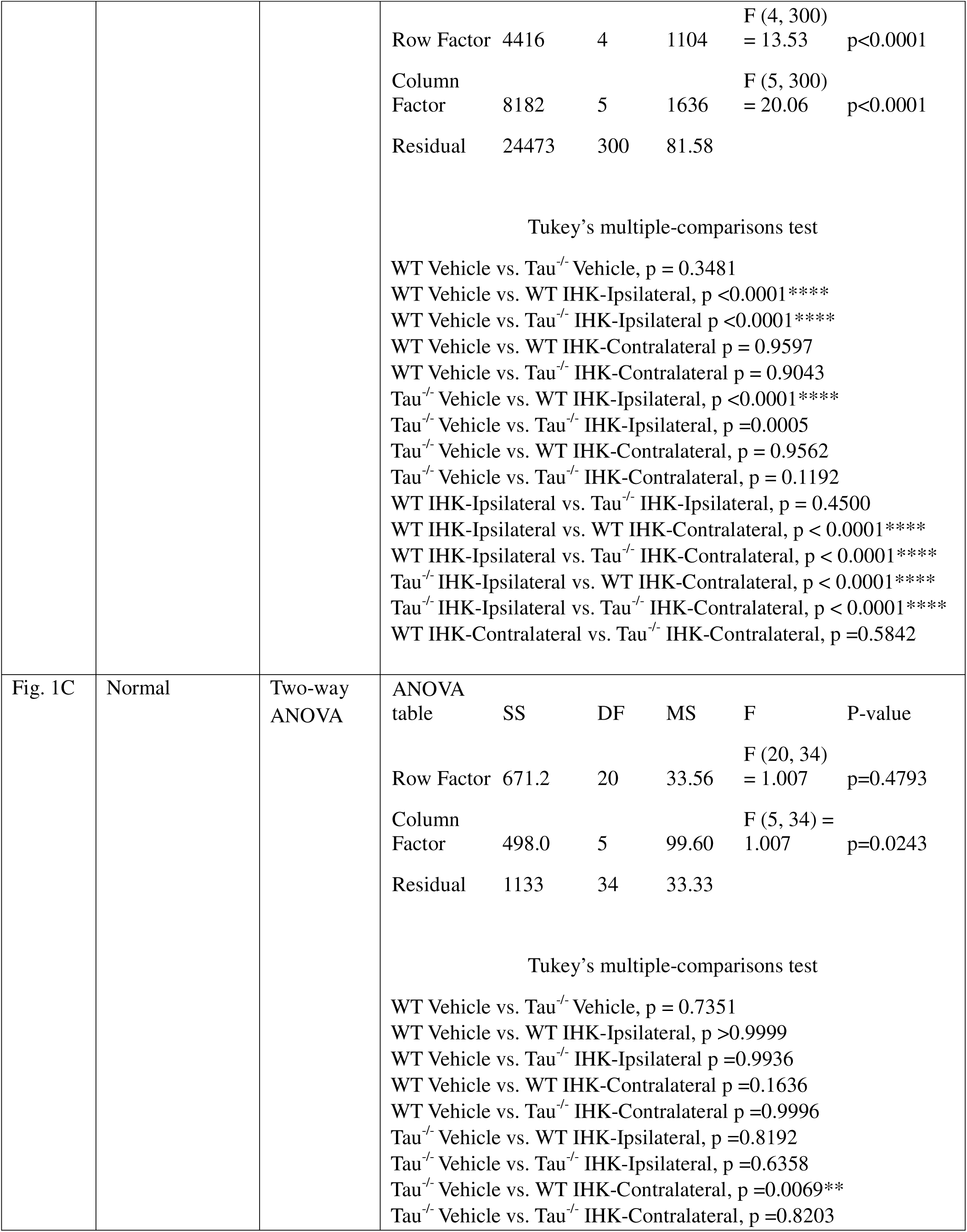
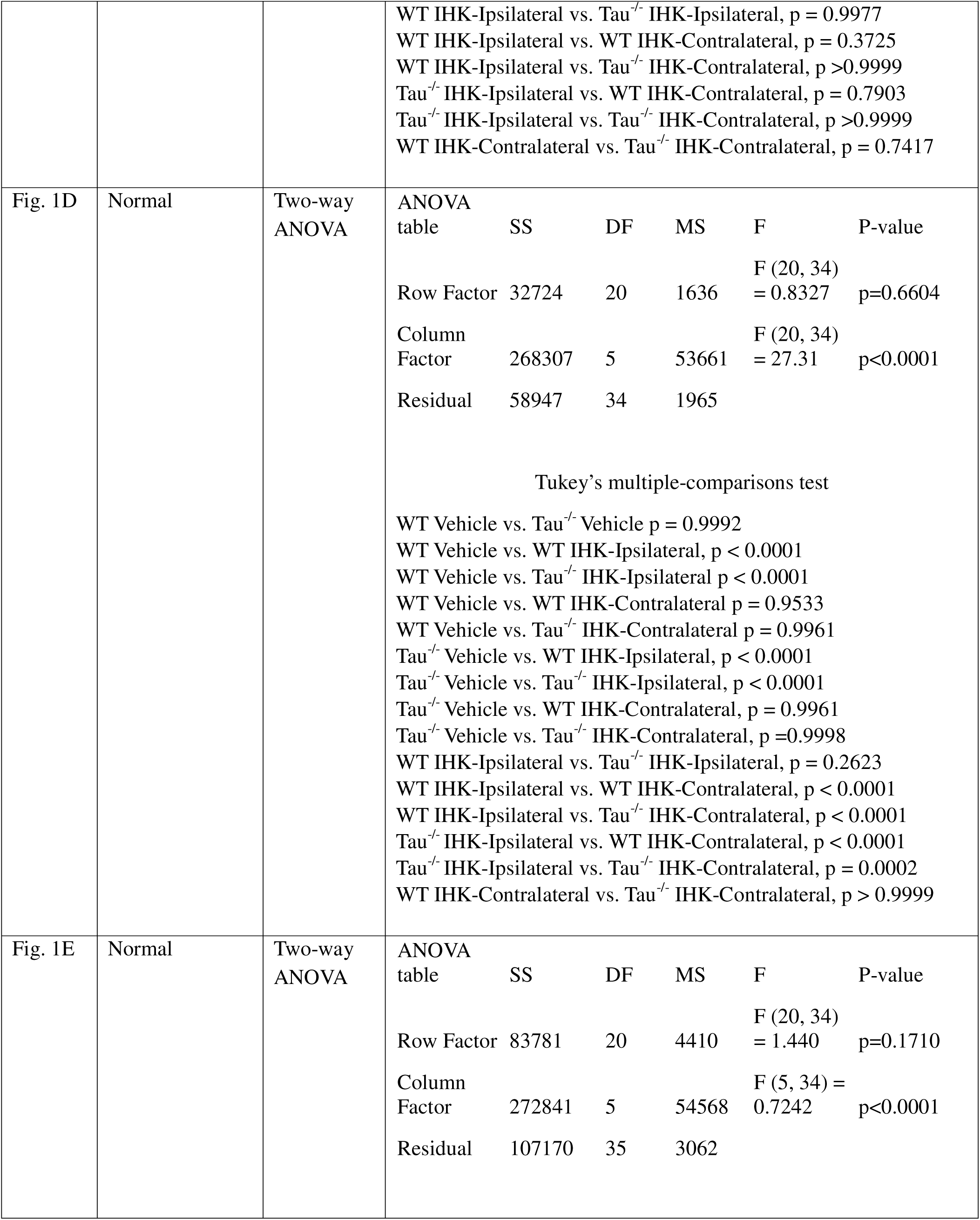
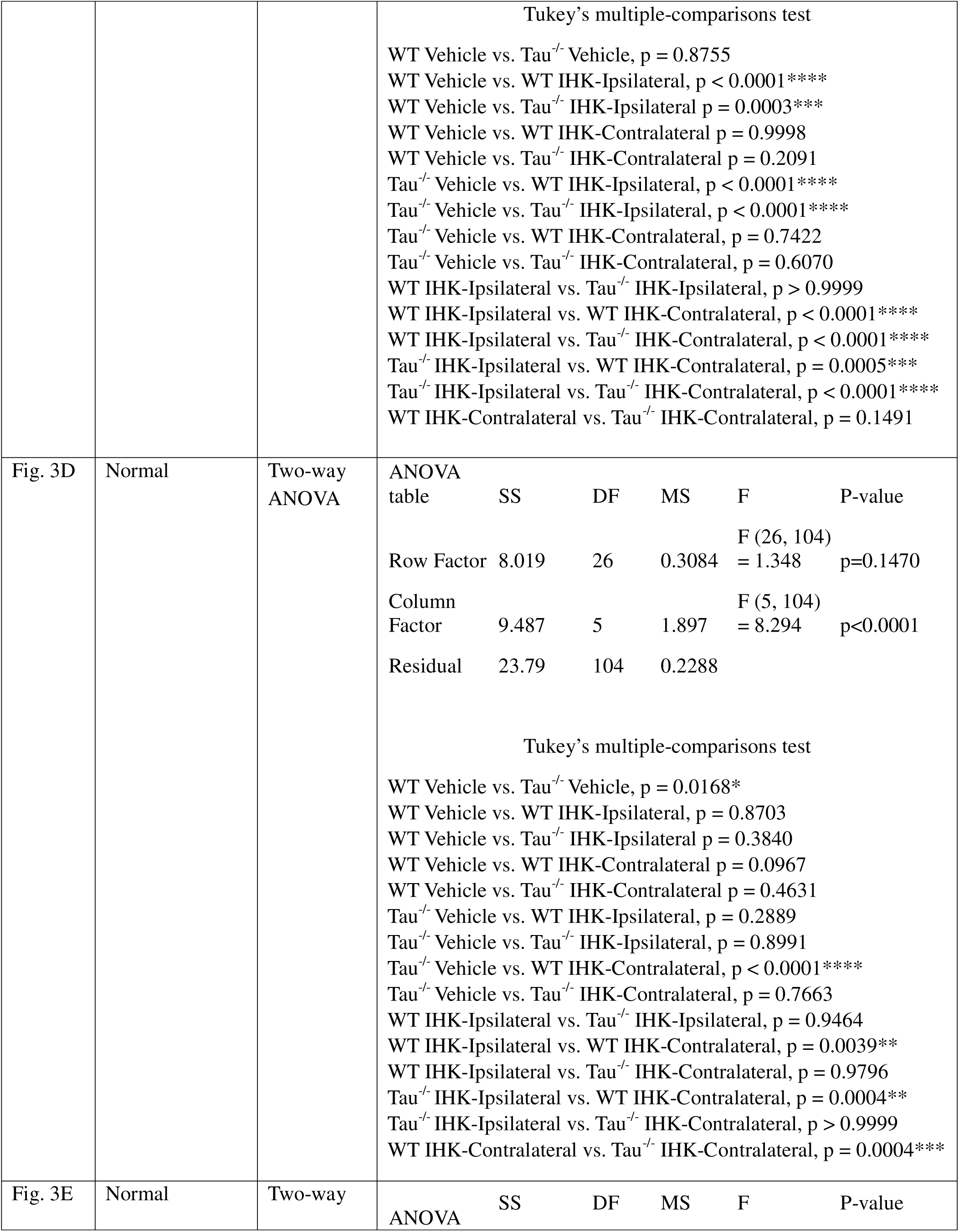
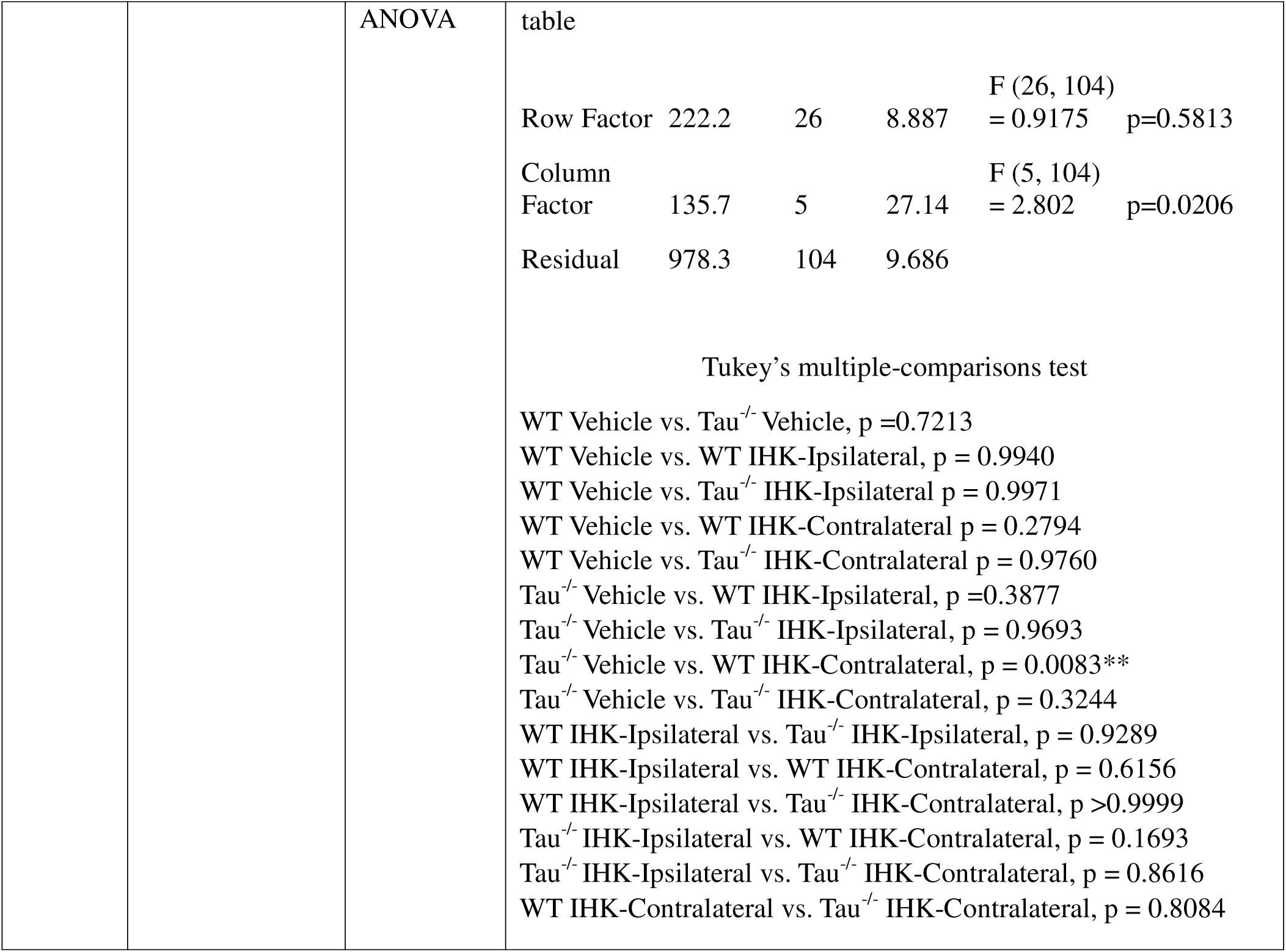
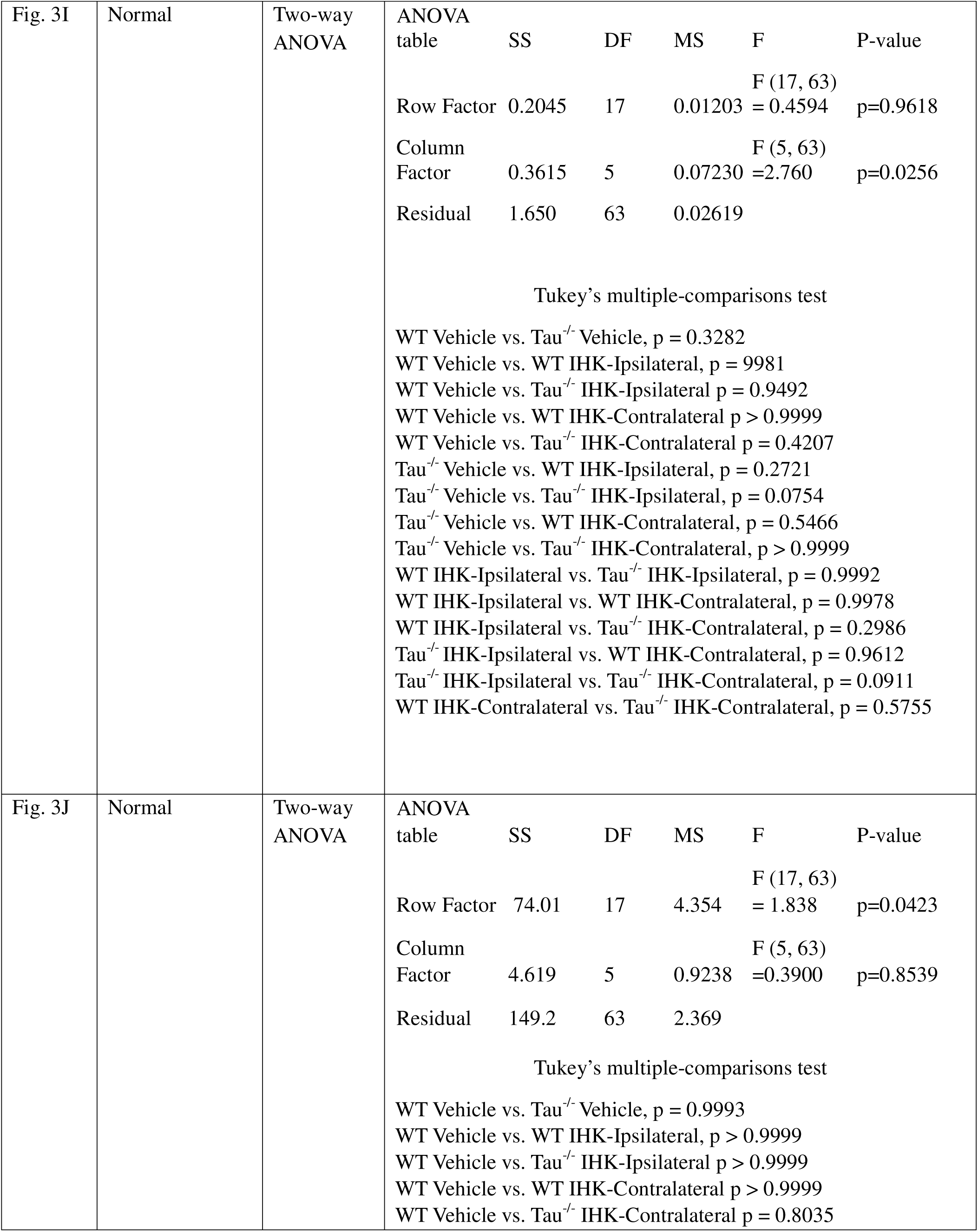
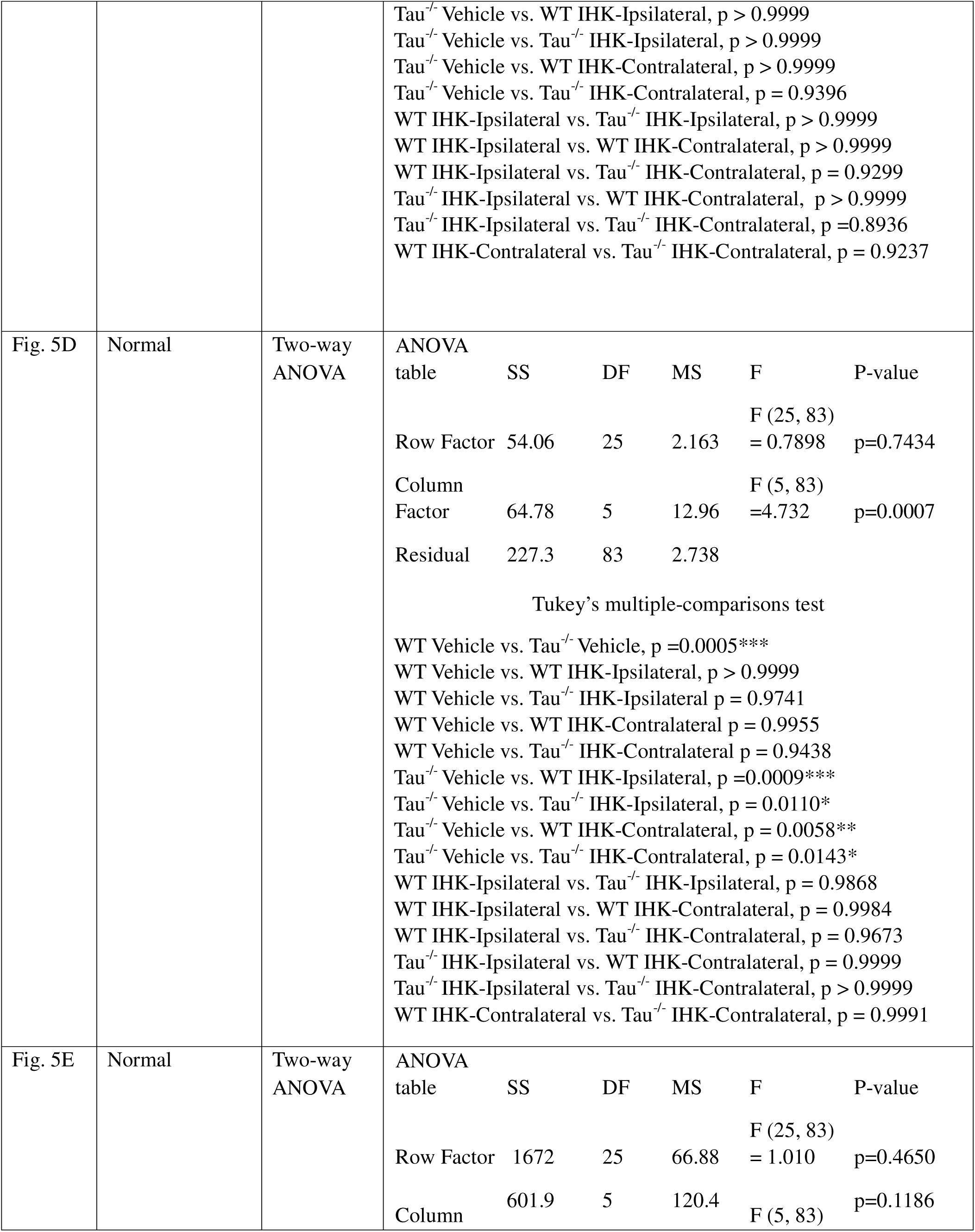
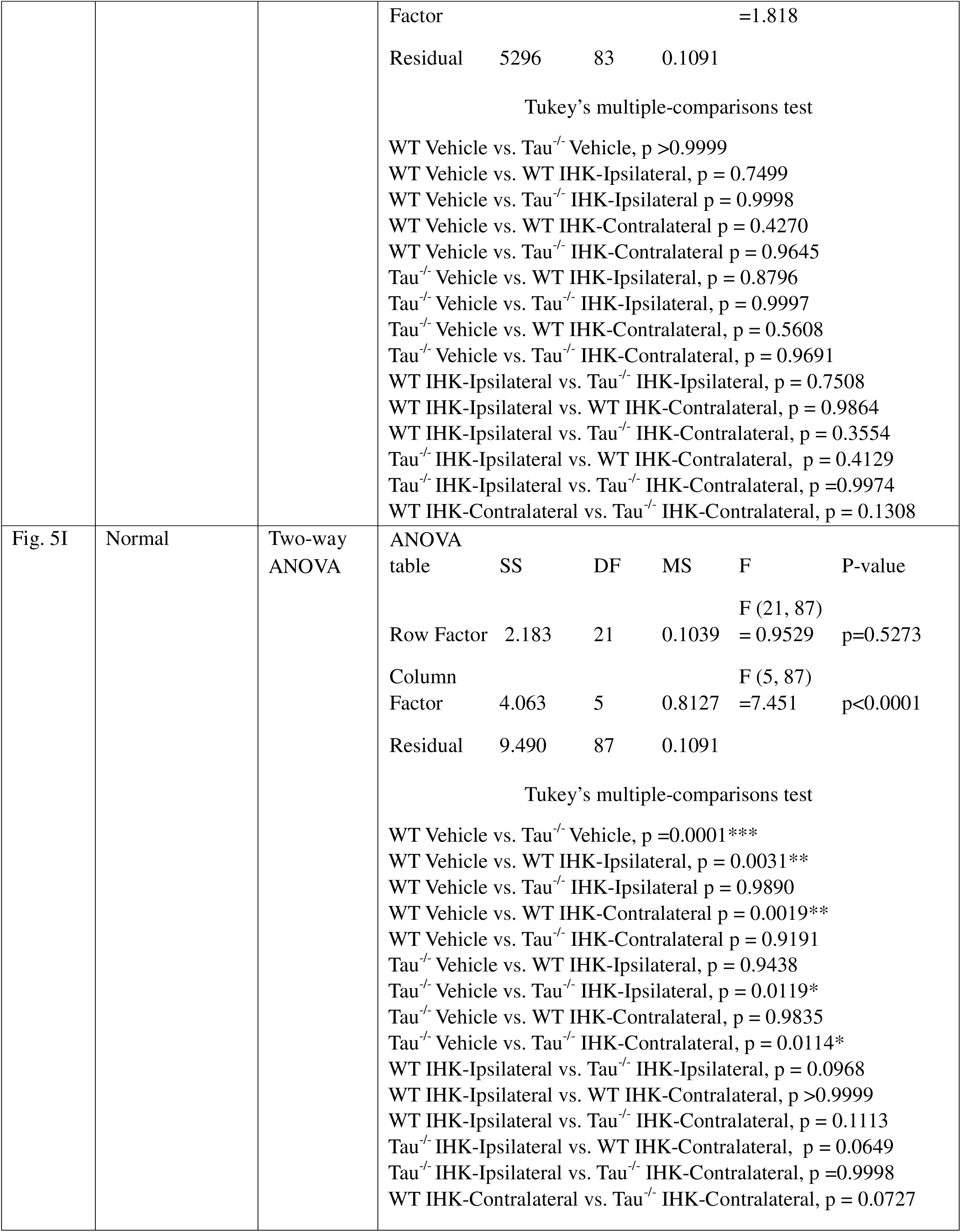
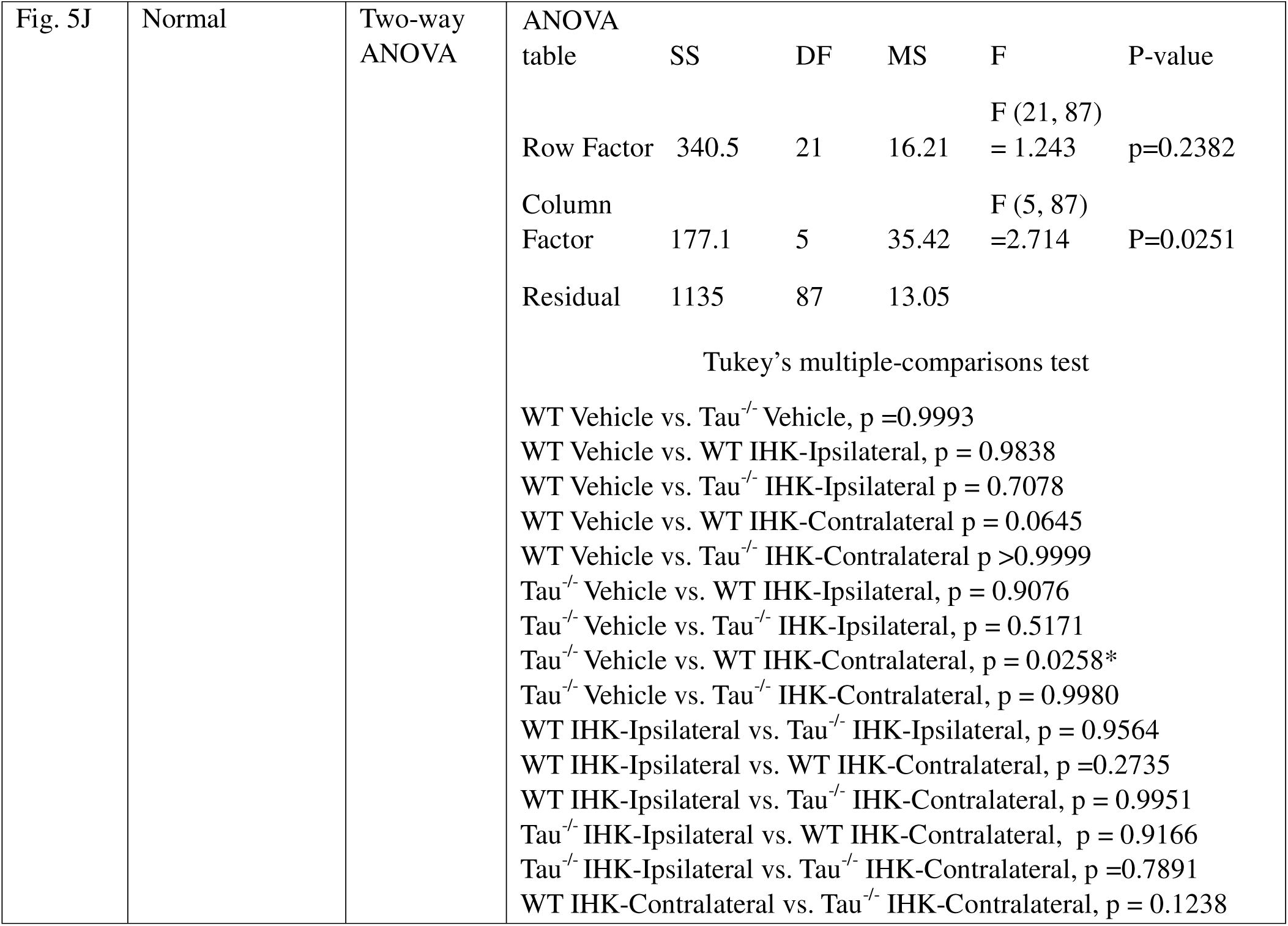
Statistical analysis for all figures.

## Results

### Intrinsic excitability of ventral DGCs increases in mice that develop TLE

In animal models of TLE and in patients (Stegen *et al*., 2012), epileptogenesis is associated with an increase in neuronal excitability of DGCs, which can be at least partly attributed to sprouting of DGC axons (i.e., mossy fiber sprouting, MFS), increased synaptic connectivity between DGCs, and reduced synaptic inhibition (Scharfman *et al*., 1999; Kobayashi & Buckmaster, 2003; Winokur *et al*., 2004; Hunt *et al*., 2010; Althaus *et al*., 2015; Lee *et al*., 2024). Differently, unilateral injection of kainate resulted in severe hippocampal damage in the dorsal hippocampus and was associated with a significant reduction in intrinsic excitability of dorsal DGCs (dDGCs) ipsilateral to the IHK injection (Fig. 1). After development of TLE, dDGCs in wildtype and tau^-/-^ mice exhibited significantly reduced action potential firing in response to depolarizing current steps, higher rheobase, and lower input resistance compared to dDGCs from vehicle-treated controls and dDGCs contralateral to the IHK injection (p<0.05; Fig. 1A2, 1A4, 1A5). Interestingly, there was no significant difference detected in the resting membrane of dDGCs across genotypes or treatments (Fig. 1A3; p>0.05). There was no difference detected in any intrinsic property between saline-treated wildtype and tau^-/-^ mice, ipsilateral dDGCs from IHK-treated wildtype and tau^-/-^ mice, or contralateral dDGCs from IHK-treated wildtype and tau^-/-^ mice (p>0.05). These results suggest DGCs in the dorsal dentate gyrus, ipsilateral to the focal kainate injection where hippocampal damage is most severe, exhibit significantly reduced excitability after TLE develops, regardless of tau expression. Since the site near the IHK injection may not reflect more widespread patterns of excitability, we examined DGCs in the ventral hippocampus to elucidate how development of TLE may alter cellular excitability and synaptic transmission of vDGCs.

To test the hypothesis that development of TLE modifies neuronal excitability of vDGCs and to determine if excitability is attenuated once TLE develops in mice lacking tau expression, we assayed intrinsic properties of vDGCs in vehicle and IHK-treated wildtype and tau^-/-^ mice that developed spontaneous seizures (Fig. 1). Action potential firing frequency in vDGCs both ipsilateral and contralateral to the IHK injection was increased in all IHK-treated mice that developed spontaneous seizures compared to vDGCs in vehicle-treated controls, with no detectable differences between genotypes (Fig. 1B2, Table 2; p<0.05). There was no significant difference in the firing frequency of vDGCs between vehicle-treated wildtype and tau^-/-^ mice (p>0.05). Interestingly, the resting membrane potential of vDGCs ipsilateral and contralateral to IHK injection in wildtype mice that developed TLE was significantly depolarized compared to vehicle-treated wildtype controls and IHK-treated tau^-/-^ mice that developed TLE (Fig. 1B3, Table 2; p<0.05). Even so, there was no significant difference detected in rheobase or input resistance across treatments or genotypes (Fig. 1B4, 1B5, Table 2; p>0.05). Together this suggests that in the ventral dentate gyrus, DGCs exhibit increased action potential firing in response to a depolarizing stimulus once epilepsy develops in the IHK model. Additionally, it is likely that tau protein plays a minimal role in regulating endogenous action potential firing of DGCs in mice of this age (14-16 wks old), consistent with a previous report (Cloyd *et al*., 2021).

**Table 2.** Summary of resting membrane potential, rheobase, and input resistance.

|  | Dorsal DGCs |  |  | Ventral DGCs |  |  |
| --- | --- | --- | --- | --- | --- | --- |
|  | RMV | Rheobase | Input | RMV | Rheobase | Input |

|  |  |  | (mV) | (pA) | Resistance<br>(mOhm) | (mV) | (pA) | Resistance<br>(mOhm) |
| --- | --- | --- | --- | --- | --- | --- | --- | --- |
| Wildtype (Ipsilateral and Contralateral) |  | Veh | -73.59 ± 1.196 (12) | 76.67 ± 9.156 (12) | 275.2 ± 13.47 (12) | -72.16 ± 0.6856 (25) | 58.33 ± 3.166 (25) | 267.4 ± 15.28 (25) |
| Tau<sup>-/-</sup> (Ipsilateral and Contralateral) |  | Veh | -74.58 ± 1.265 (21) | 78.00 ± 7.157 (21) | 276.3 ± 14.45 (21) | -70.87 ± 1.784 (20) | 58.00 ± 4.078 (20) | 311.4 ± 21.02 (20) |
| Wildtype | Ipsilateral | IHK | -72.92 ± 0.7083 (5) | 296.0 ± 24.00 (5)*<sup>#</sup> | 119.7 ± 6.097 (5)*<sup>#</sup> | -65.34 ± 1.890 (19)* | 55.79 ± 3.613 (19) | 237.6 ± 30.47 (19) |
|  | Contralateral | IHK | -67.44 ± 2.610 (11) | 61.82 ± 9.892 (11)<sup>+</sup> | 269.20 ± 23.53 (11)<sup>+</sup> | -66.16 ± 2.026 (20)* | 47.367 ± 3.138 (20) | 280.3 ± 18.78 (20) |
| Tau<sup>-/-</sup> | Ipsilateral | IHK | -71.13 ± 1.418 (4) | 230.0 ± 57.45 (4)<sup>**#</sup> | 110.46 ± 17.24 (4)<sup>**#</sup> | -70.38 ± 1.087 (18) | 53.33 ± 3.234 (18) | 276.5 ± 25.40 (18) |
|  | Contralateral | IHK | -72.47 ± 3.317 (4) | 70.00 ± 10.00 (4)<sup>+</sup> | 346.3 ± 24.26 (4)<sup>+</sup> | -68.91 ± 1.007 (22) | 57.27 ± 4.012 (22) | 302.9 ± 24.79 (22) |

\* different from vehicle wildtype;
\*\* different from vehicle tau<sup>-/-</sup>;
<sup>+</sup> different from ipsilateral same genotype;
<sup>#</sup> different from contralateral same genotype

### Development of TLE in tau^-/-^ mice does not lead to excitatory synaptic reorganization in the ventral dentate gyrus

We recently showed that the development of spontaneous seizures coincides with significantly increased excitatory synaptic input to dDGCs ipsilateral to the IHK lesion in both wildtype and tau^-/-^ mice (Moseley *et al*., 2025). We hypothesized that seizure expression or epileptogenesis is modified in mice lacking tau expression due to suppressed excitatory synaptic reorganization distant from the kainate lesion. As an assay of overall excitatory synaptic input (action potential dependent and independent), we examined the frequency of sEPSCs in vDGCs from tau^-/-^ and wildtype mice that developed TLE. In wildtype mice that developed spontaneous seizures, significantly increased sEPSC frequency was observed in vDGCs contralateral to the IHK injection compared to vDGCs in the ipsilateral hemisphere (Fig. 2D, Table 3; p<0.05). However, no significant differences were detected between vDGCs from vehicle- and IHK-treated wildtype mice (Fig. 2D, Table 3; p>0.05). Development of TLE in mice lacking tau expression did not correspond with differences in sEPSC frequency (Fig. 2D, Table 3; p>0.05). Interestingly, we found that sEPSC frequency in vDGCs from naive, vehicle-treated tau^-/-^ mice was significantly lower than in wildtype controls (Fig 2D, Table 3; p<0.05), suggesting tau expression may play a role in excitatory synaptic transmission in the local dentate gyrus network. As in the dorsal dentate gyrus (Moseley *et al*., 2025), differences in the amplitude of unitary sEPSC events were not detected in the vDGCs across all mice (Fig 2E, Table 3; p>0.05). Thus, changes in sEPSC frequency in vDGCs was a feature of epileptogenesis in wildtype mice that was not conserved in tau^-/-^ mice.

**Figure 2.**
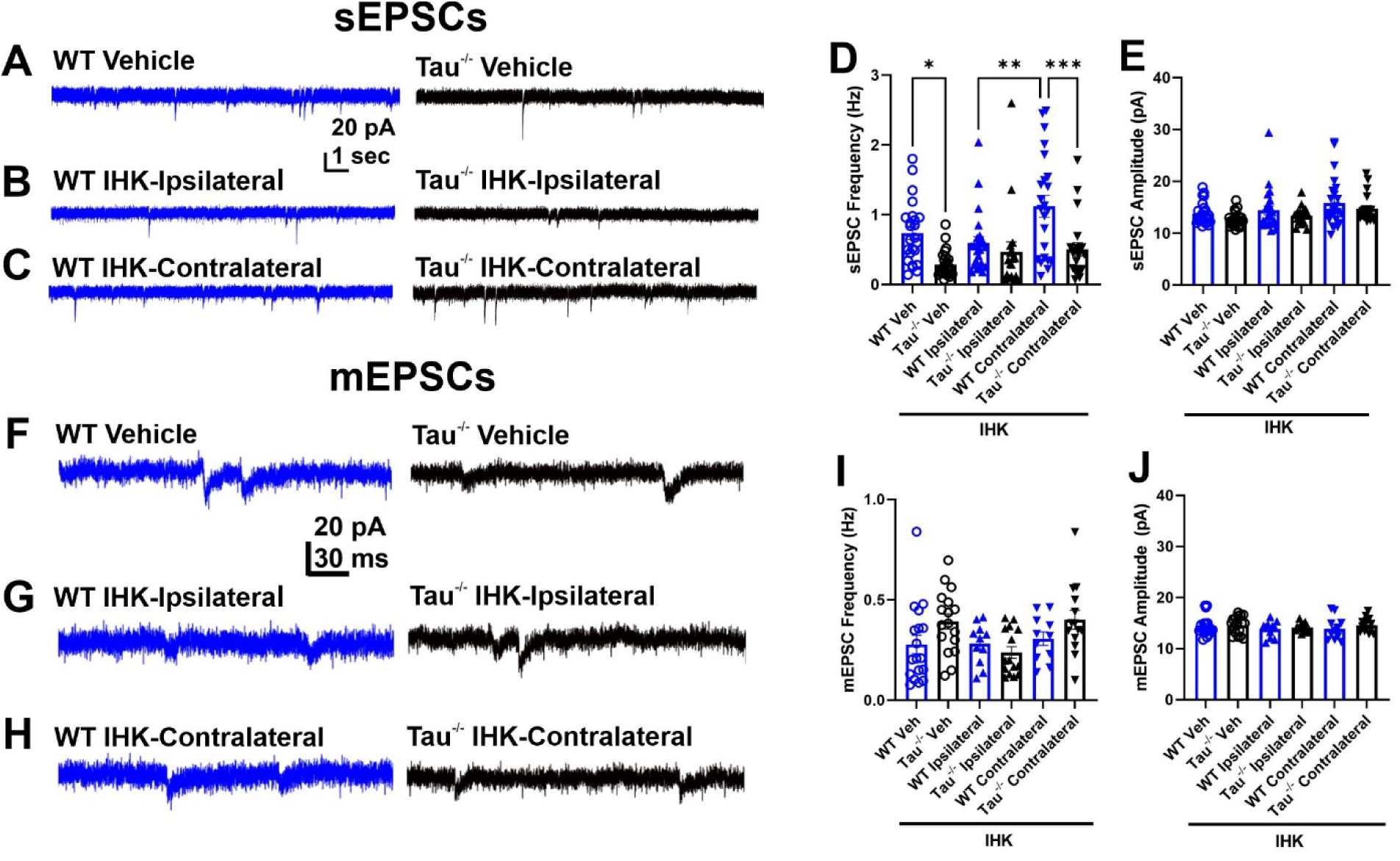
Synaptic excitation to vDGCs is not increased after development of TLE in mice lacking tau expression. (A) Representative traces showing sEPSCs from vehicle-treated WT and tau^-/-^ mice. (B) Representative sEPSC traces from vDGCs ipsilateral to IHK injection in IHK-treated WT and tau^-/-^ mice. (C) Representative sEPSC traces from vDGCs contralateral to IHK injection in IHK-treated WT and tau^-/-^ mice. (D) sEPSC frequency from vehicle and IHK-treated wildtype and tau^-/-^ mice (*p<0.05, **p<0.01, ***p<0.001, Two-way ANOVA). (E) Amplitude of sEPSC events from vehicle and IHK-treated wildtype and tau^-/-^ mice (p>0.05, Two-way ANOVA). WT Vehicle: N=7, n=25; Tau^-/-^ Vehicle: N=8, n=25; WT IHK: N=10, ipsilateral: n=23, contralateral: n=23; Tau^-/-^ IHK N=10, ipsilateral: n=18, contralateral: n=21). (F) Representative mEPSC traces from vehicle-treated wildtype and tau^-/-^ mice. (G) Representative mEPSCs traces from vDGCs ipsilateral and contralateral to IHK injection in IHK-treated wildtype mice. (H) Representative mEPSC traces from vDGCs ipsilateral and contralateral to IHK injection in IHK-treated tau^-/-^ mice. (I) mEPSC frequency from vehicle and IHK-treated wildtype and tau^-/-^ mice (p>0.05, Two-way ANOVA). (J) Amplitude of mEPSC events from vehicle and IHK-treated wildtype and tau^-/-^ mice (p>0.05, Two-way ANOVA). Traces from WT mice are blue; from tau^-/-^ are black for display purposes. WT Vehicle: N=4, n=18; Tau^-/-^ Vehicle: N=4, n=17; WT IHK: N=4, ipsilateral: n=11, contralateral: n=11; Tau^-/-^ IHK N=3, ipsilateral: n=15, contralateral: n=14).

**Table 3.** Summary of voltage-clamp electrophysiology measurements.

| Treatment and Genotype |  |  | Point Estimate ± SEM (n) |  |  |  |  |  |  |  |
| --- | --- | --- | --- | --- | --- | --- | --- | --- | --- | --- |
|  |  |  | sEPSC<br>Frequency<br>(Hz) | sEPSC<br>Amplitude<br>(pA) | sIPSC<br>Frequency<br>(Hz) | sIPSC<br>Amplitude<br>(pA) | mEPSC<br>Frequency<br>(Hz) | mEPSC<br>Amplitude<br>(pA) | mIPSC<br>Frequency<br>(Hz) | mIPSC<br>Amplitude<br>(pA) |
| Wildtype (Ipsilateral and Contralateral) |  | Veh | 0.7375 ± 0.0905 (25) | 13.84 ± 0.4340 (25) | 3.579 ± 0.2226 (24) | 29.83 ± 1.360 (24) | 0.2777 ± 0.046 (18) | 14.44 ± 0.4665 (18) | 1.923 ± 0.0903 (22) | 20.41 ± 0.9307 (22) |
| Tau<sup>-/-</sup> (Ipsilateral and Contralateral) |  | Veh | 0.2833 ± 0.0351 (27)* | 12.56 ± 0.2539 (27) | 1.175 ± 0.1288 (15)* | 29.83 ± 1.294 (15) | 0.3923 ± 0.038 (17) | 14.57 ± 0.4185 (17) | 0.7141 ± 0.0577 (21)* | 20.12 ± 0.8074 (21) |
| Wildtype | Ipsilateral | IHK | 0.5930 ± 0.0909 (23)<sup>#</sup> | 14.41 ± 0.8492 (23) | 3.482 ± 0.3475 (25) | 33.63 ± 2.014 (25) | 0.2813 ± 0.0306 (11) | 13.80 ± 0.4747 (11) | 0.8066 ± 0.0626 (21)* | 21.27 ± 0.7384 (21) |
|  | Contralateral | IHK | 1.1213 ± 0.1543 (23)<sup>+</sup> | 15.86 ± 1.005 (23) | 3.498 ± 0.5692 (17) | 35.44 ± 2.102 (17) | 0.3064 ± 0.0336 (11) | 13.89 ± 0.2125 (11) | 0.7995 ± 0.0491 (20)* | 23.79 ± 0.7460 (20) |
| Tau<sup>-/-</sup> | Ipsilateral | IHK | 0.4662 ± 0.1451 (18) | 13.35 ± 0.4249 (18) | 3.263 ± 0.4572 (16)<sup>+</sup> | 29.10 ± 2.140 (16) | 0.2382 ± 0.0293 (15) | 14.16 ± 0.2125 (15) | 1.158 ± 0.1085 (13)<sup>+</sup> | 22.30 ± 1.240 (13) |
|  | Contralateral | IHK | 0.5038 ± 0.0945 (21)<sup>b</sup> | 14.71 ± 0.6170 (21) | 3.181 ± 0.3824 (17)<sup>#</sup> | 28.05 ± 2.141 (17) | 0.4010 ± 0.0473 (14) | 14.60 ± 0.3630 (14) | 1.130 ± 0.0879 (17)<sup>#</sup> | 20.38 ± 0.6688 (17) |

\* different from vehicle wildtype;
\*\* different from vehicle tau<sup>-/-</sup>;
<sup>+</sup> different from ipsilateral same genotype;
<sup>#</sup> different from contralateral same genotype;
<sup>a</sup> different from ipsilateral WT-IHK
<sup>b</sup> different from contralateral WT-IHK

To determine if the changes in excitatory synaptic input to vDGCs in the wildtype mice that developed TLE resulted from action potential dependent or independent release of glutamate onto ventral DGCs, we assayed mEPSCs in the presence of TTX. In all mice, regardless of treatment or genotype, no significant differences in mEPSC frequency or amplitude were observed in vDGCs from mice with TLE (Fig. 2I-J; Table 3; p>0.05). Furthermore, unlike sEPSCs, there was no significant difference in mEPSC frequency between naïve vehicle-treated tau^-/-^ mice and wildtype controls, suggesting that lack of tau expression reduces action-potential dependent glutamate release to vDGCs.

Spontaneous seizure development in rodent TLE models and in patients with TLE has been shown to be associated with excitatory synaptic reorganization that is supported by MFS into the inner molecular layer of the dentate gyrus (Franck *et al*., 1995). It is hypothesized that the sprouting of DGC axons onto other DGCs contributes to recurrent excitation in the circuit leading to a lower spontaneous seizure threshold (Buckmaster & Dudek, 1997). While it is debated whether the severity of MFS is associated with seizure burden, it is relatively unknown if modified epileptogenesis contributes to varying degrees of MFS. Mice lacking tau expression develop TLE but at a lower rate with a reduced severity of spontaneous seizures than wildtype controls. In several rodent models of TLE, it has been shown that MFS is detectable in the ventral dentate gyrus once epilepsy develops (Zeidler *et al*., 2018). Therefore, we investigated whether tau^-/-^ mice exhibit MFS in the ventral dentate gyrus associated with development of TLE after focal injection of kainate in the dorsal hippocampus. To visualize potential MFS, we used Timm’s stain that allows for detection of zinc containing axons of DGCs. Timm’s scoring was quantified as described (Cavazos *et al*., 1991). The presence and distribution of Timm positive granules in the molecular layer was scored on a scale of 0-5 (0: no granules, 1: sparse granule distribution, 2: continuous granule distribution, 3: prominent continuous granules, 4: granules forming a dense laminar band in the molecular layer, 5: confluent dense laminar band extending farther into the molecular layers). After development of TLE, wildtype mice (N=5) and tau^-/-^ mice (N=4) exhibited MFS in the inner molecular layer both ipsilateral and contralateral to IHK injection (Fig. 3). Semi-quantitative analysis of Timm’s score did not reveal any detectable differences in the severity of MFS in the ventral dentate gyrus between IHK-treated wildtype and tau^-/-^ mice (Fig. 3D). Notably, MFS was not observed in vehicle treated mice of either genotype (wildtype: N=2; tau^-/-^ N=2) or in IHK-treated wildtype or tau^-/-^ mice that did not exhibit spontaneous seizures during the video recording period (WT: N=3, Tau^-/-^ N=1). Therefore, development of TLE in the IHK model results in MFS in the ventral dentate gyrus, distant from the focal kainate injection in the dorsal hippocampus in both genotypes, even though evidence of increased glutamate release was only observed in wildtype mice. Increased excitatory synaptic input to vDGCs may therefore not entirely correlate with the presence of MFS.

**Figure 3.**
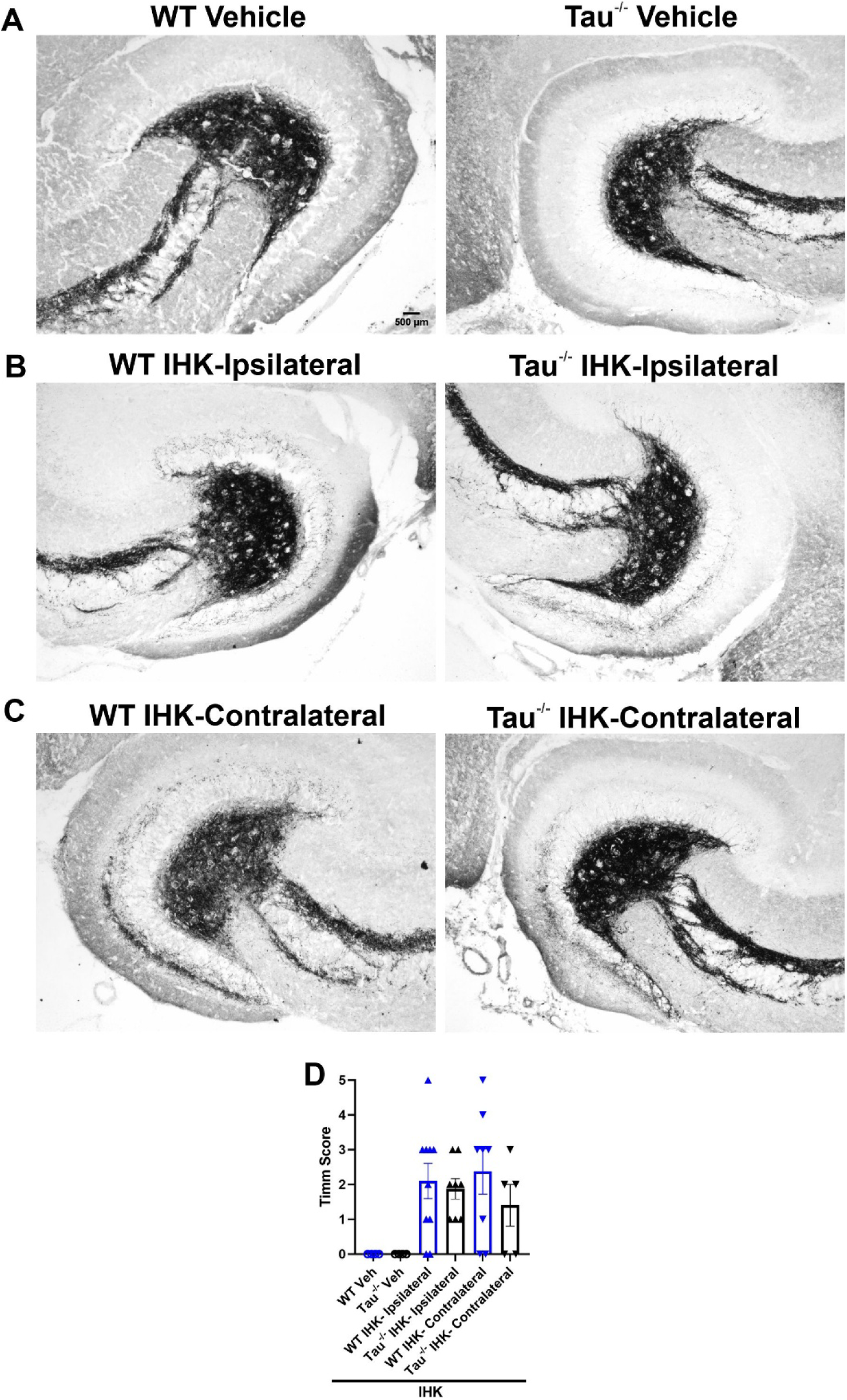
After development of TLE mossy fiber sprouting is detected in the ventral dentate gyrus inner molecular layer of IHK treated mice in both genotypes. (A) Representative Timm’s stain in the ventral dentate gyrus from vehicle-treated WT and tau^-/-^ mice (WT: N=2, Tau^-/-^: N=2). (B) Representative Timm’s stain in the ventral dentate gyrus from IHK-treated WT mice (WT: N=5). (C) Representative Timm’s stain in the ventral dentate gyrus from IHK-treated tau^-/-^ mice (Tau^-/-^: N=5). (D) Semi-quantittive assessment of Timm score of ventral dentate gyrus sections from vehicle and IHK-treated WT and tau^-/-^ mice. 0=no granules, 1=sparse granules in the supragranular layer, 2=more granules in a continuous distribution, 3=prominent granules in the supragranular layer, 4=prominent granules in the supragranular layer forming a dense band, 5=confluent laminar band in the supragranular layer. Mossy fiber sprouting (MFS) was detected after IHK in both WT and tau^-/-^ mice while no MFS was observed in vehicle treated controls.

### Ventral DGCs in wildtype mice receive less inhibitory synaptic input than in tau^-/-^ mice but development of TLE increases synaptic inhibition in mice lacking tau expression

We previously demonstrated that development of TLE in tau^-/-^ mice results in increased inhibitory synaptic input to dorsal DGCs contralateral to the IHK injection compared to vehicle-treated controls; this increase was not observed in wildtype mice (Moseley *et al*., 2025). We investigated whether similar inhibitory synaptic reorganization was detectable at more distal sites in the ventral dentate gyrus. We recorded sIPSCs in vDGCs as an assay for overall inhibitory synaptic input (Fig. 4). Unlike for sEPSC frequency in vDGCs (Table 3), and unlike for sIPSC frequency in dDGCs reported previously (Moseley *et al*., 2025), there was no difference in sIPSC frequency in vDGCs from wildtype mice that developed TLE compared to vehicle-treated controls (Fig. 4D, Table 3; p>0.05), suggesting that inhibitory synaptic input remains intact in the ventral dentate gyrus distant from the focal injury in the dorsal hippocampus. Interestingly, sIPSC frequency was significantly increased both ipsilateral and contralateral to the IHK injection in tau^-/-^ mice that developed TLE compared to vehicle treated tau^-/-^ mice (Fig. 4D, Table 3; p<0.05), suggesting inhibitory synaptic reorganization previously seen in the dorsal contralateral hemisphere in tau^-/-^ mice occurs throughout the dentate gyrus in these mice. Additionally, like sEPSC frequency, sIPSC frequency was also significantly lower in vehicle-treated tau^-/-^ mice compared to wildtype controls (Fig. 4D, Table 3; p<0.05), like what we observed in the dorsal dentate gyrus (Moseley *et al*., 2025) and consistent with a role of tau in development of inhibitory circuitry. There was no detectable difference in sIPSC event amplitude across genotypes or treatments (Fig. 4E, Table 3; p>0.05).

**Figure 4.**
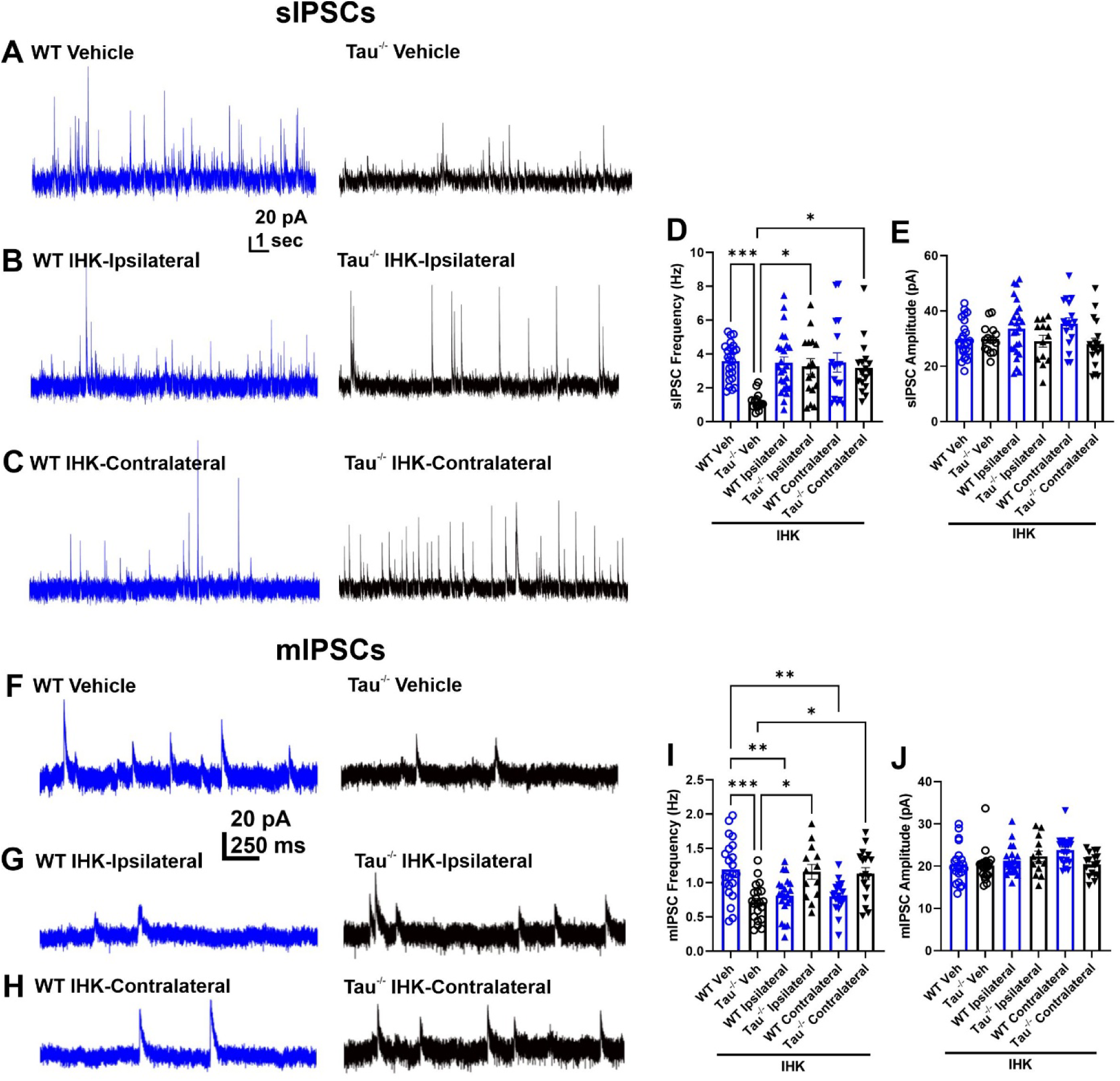
Ventral DGCs in mice lacking tau expression receive less inhibitory synaptic input and development of TLE increases inhibitory input in tau^-/-^ but not WT mice. (A) Representative traces showing sIPSC from vehicle-treated wildtype (WT) and tau^-/-^ mice. (B) Representative sIPSC traces from vDGCs ipsilateral to IHK injection in IHK-treated WT and tau^-/-^ mice. (C) Representative sIPSC traces from vDGCs contralateral to IHK injection in IHK-treated WT and tau^-/-^ mice. (D) sIPSC frequency from vehicle and IHK-treated wildtype and tau^-/-^ mice (*p<0.05, ***p<0.001, Two-way ANOVA). (E) Amplitude of sIPSC events from vehicle and IHK-treated wildtype and tau^-/-^ mice (p>0.05, Two-way ANOVA). WT Vehicle: N=7, n=24; Tau^-/-^ Vehicle: N=8, n=15; WT IHK: N=10, ipsilateral: n=25, contralateral: n=17 ; Tau^-/-^ IHK N=10, ipsilateral: n=16, contralateral: n=17). (F) Representative mIPSC traces from vehicle-treated wildtype and tau^-/-^ mice. (G) Representative mIPSCs traces from vDGCs ipsilateral and contralateral to IHK injection in IHK-treated wildtype mice. (H) Representative mIPSC traces from vDGCs ipsilateral and contralateral to IHK injection in IHK-treated tau^-/-^ mice. (I) mIPSC frequency from vehicle and IHK-treated wildtype and tau^-/-^ mice (*p<0.05, **p<0.01, ***p<0.001, Two-way ANOVA). (J) Amplitude of mEPSC events from vehicle and IHK-treated wildtype and tau^-/-^ mice (p>0.05, Two-way ANOVA). WT Vehicle: N=3, n=18; Tau^-/-^ Vehicle: N=4, n=17; WT IHK: N=7, ipsilateral: n=11, contralateral: n=11; Tau^-/-^ IHK N=5, ipsilateral: n=15, contralateral: n=14). Traces from WT mice are blue and from tau^-/-^mice are black for clarity.

To assess action potential independent inhibitory synaptic events, we recorded mIPSCs in the presence of TTX (Fig. 4). After development of TLE, mIPSC frequency was significantly lower both ipsilateral and contralateral to the IHK injection in wildtype mice compared to vehicle treated controls (Fig. 4I, Table 3; p<0.05). Like sIPSC frequency, mIPSC frequency was significantly increased in tau^-/-^ mice that developed TLE compared to vehicle-treated tau^-/-^ mice (Fig. 4I, Table 3; p<0.05). Together this suggests that after the development of TLE, there is a potential loss of functional inhibitory synapses and/or reduced vesicular release of GABA onto vDGCs in mice that express tau (i.e., wildtype mice), whereas development of TLE in mice lacking tau expression results in increased inhibitory synaptic transmission bilaterally and distant to focal injury.

Interestingly, like sIPSC frequency, mIPSC frequency was significantly lower in vehicle-treated tau^-/-^ mice compared to wildtype controls (Fig. 4I, Table 3; p<0.05). There was no significant difference detected in mIPSC event amplitude between IHK-treated mice of either genotype that developed TLE compared to their vehicle-treated controls (p>0.05), consistent with preserved postsynaptic GABA receptor function in vDGCs even after epilepsy develops (Fig. 4J, Table 3). This highlights a potential mechanism of tau protein’s endogenous function in inhibitory synaptic transmission in which lack of tau expression may suppress the function of GABAergic synapses normally and contribute to epileptogenesis, whereas TLE development increases functional interneuron connectivity with DGCs in the local dentate gyrus network, possibly in a compensatory manner, once epilepsy develops in tau^-/-^ mice.

### Development of TLE does not alter excitatory synaptic input to inhibitory interneurons in the ventral dentate gyrus

Development of TLE in mice lacking tau expression is associated with increased inhibitory synaptic input to both ventral and dorsal (Moseley *et al*., 2025) DGCs. In several rodent models of TLE, inhibitory interneurons can receive increased excitatory input from sprouted synapses or boutons arising from DGCs (Buckmaster & Jongen-Relo, 1999; McNeill *et al*., 2003; White *et al*., 2004; Halabisky *et al*., 2010; Kang *et al*., 2022). Therefore, we tested the hypothesis that recurrent excitation to inhibitory interneurons results in a network driven increase in inhibitory synaptic input to vDGCs after development of epilepsy in mice lacking tau expression.

Using AAV-Dlx-ChR2-mCherry expression to identify GABAergic interneurons, we recorded sEPSCs in mCherry labelled cells (Fig.5). Interestingly, our assay of sEPSCs in wildtype and tau^-/-^ mice demonstrated no significant genotype or treatment related differences in sEPSC frequency or amplitude in interneurons (p>0.05; Fig. 5D-E). This suggests that neither tau expression nor the development of epilepsy modifies overall excitatory synaptic input to interneurons, suggesting that tau protein does not play a major role in regulating excitatory synaptic transmission from DGCs in this circuit, inconsistent with what previous studies have suggested when tau expression is deleted in excitatory neurons globally (Shao *et al*., 2022). Therefore, it is likely that the modified inhibitory synaptic input to vDGCs after development of TLE is due to changes in the nature or number of the GABAergic synapses onto vDGCs, rather than excitatory input upstream from either vDGCs or surviving mossy cells. Even so, future studies are required to determine if increased excitatory synaptic input to specific interneuron subtypes occurs.

**Figure 5:**
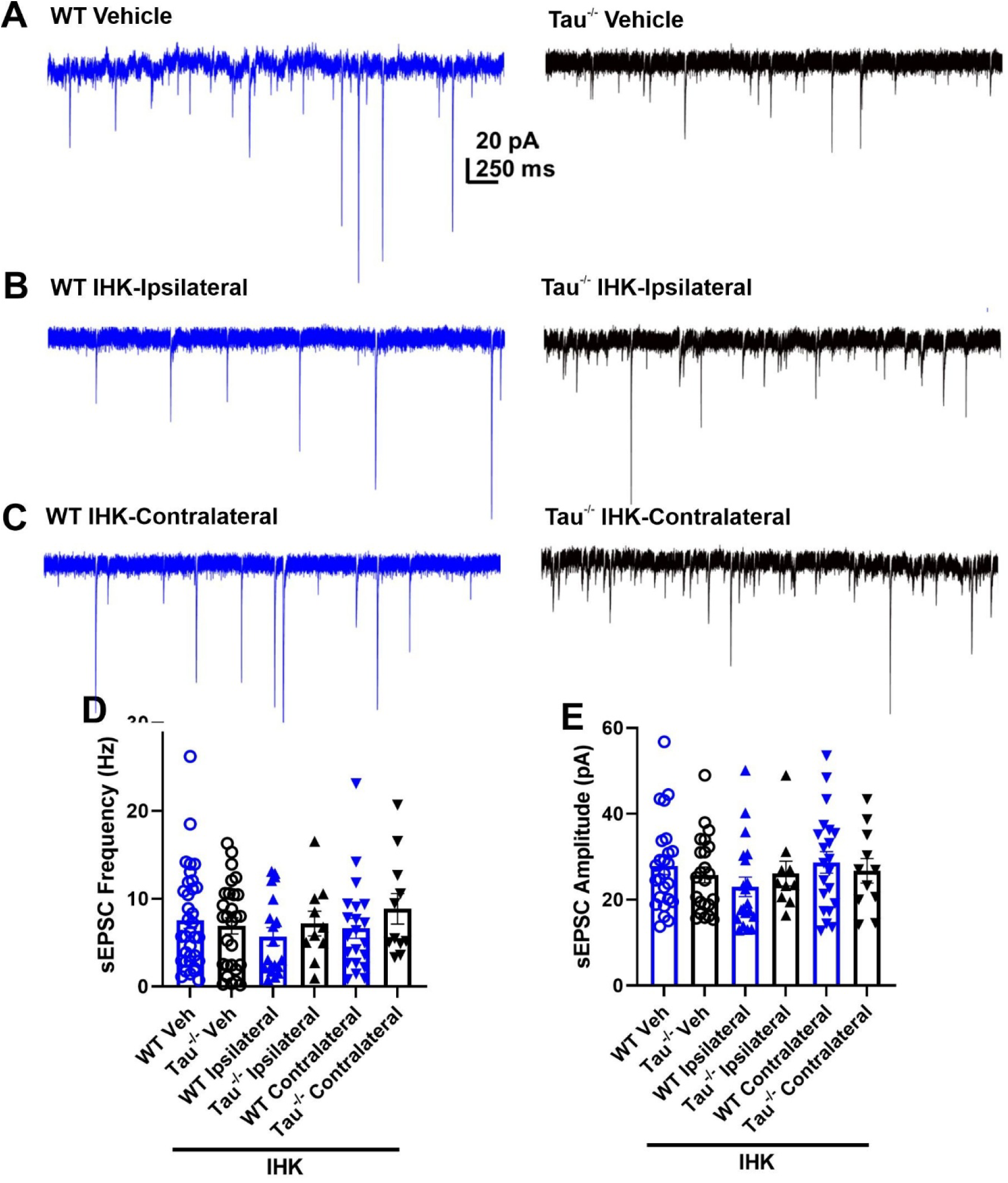
Development of TLE does not alter excitatory synaptic input to inhibitory interneurons in the ventral dentate gyrus. (A) Representative traces showing sEPSCs recorded in interneurons from vehicle-treated wildtype (WT) and tau^-/-^ mice. (B) Representative sEPSC traces from interneurons ipsilateral to IHK injection in IHK-treated WT and tau^-/-^ mice. (C) Representative sEPSC traces from interneurons contralateral to IHK injection in IHK-treated WT and tau^-/-^ mice. (D) sEPSC frequency from vehicle and IHK-treated wildtype and tau^-/-^ mice (p>0.05, Two-way ANOVA). (E) Amplitude of sEPSC events from vehicle and IHK-treated wildtype and tau^-/-^ mice (p>0.05, Two-way ANOVA). WT Vehicle: N=9, n=35; Tau^-/-^ Vehicle: N=7, n=26; WT IHK: N=5, ipsilateral: n=19, contralateral: n=21 ; Tau^-/-^ IHK N=6, ipsilateral: n=10, contralateral: n=11).

### Development of TLE increases evoked synaptic inhibition and synaptic connections between inhibitory interneurons and vDGCs in mice lacking tau expression but not in wildtype mice

Excitatory synaptic input to inhibitory interneurons appears to remain unchanged after development of epilepsy, suggesting altered inhibition in the dentate gyrus is localized to GABAergic synapses at sites distal to the kainate injection after spontaneous seizures develop. Mice expressing murine tau (i.e., wildtype mice) exhibit reduced action potential independent inhibitory synaptic events while overall inhibitory input is unchanged, potentially masked by normalized excitability of surviving interneurons. We took advantage of the ChR2 fusion to mCherry expressing interneurons to selectively activate interneurons and record evoked inhibitory post-synaptic currents (eIPSCs) in vDGCs.

Optogenetic stimulation of inhibitory interneurons expressing ChR2 (10 ms duration, 5 stimulations, 1 sec interstimulus interval) resulted in no significant differences in amplitude or charge transfer of eIPSCs recorded ipsilateral or contralateral to IHK injection in wildtype mice compared to vehicle treated controls (Fig. 6A-E; p>0.05). However, there was a significant increase in the decay time constant of eIPSCs contralateral to IHK injection compared to vehicle controls (Fig. 6F; p<0.05). Further, no significant difference in the number of successful connections between stimulated interneurons and vDGCs ipsilateral or contralateral in IHK treated wildtype mice compared to vehicle controls was detected (Fig. 6G-J; p>0.05).

**Figure 6.**
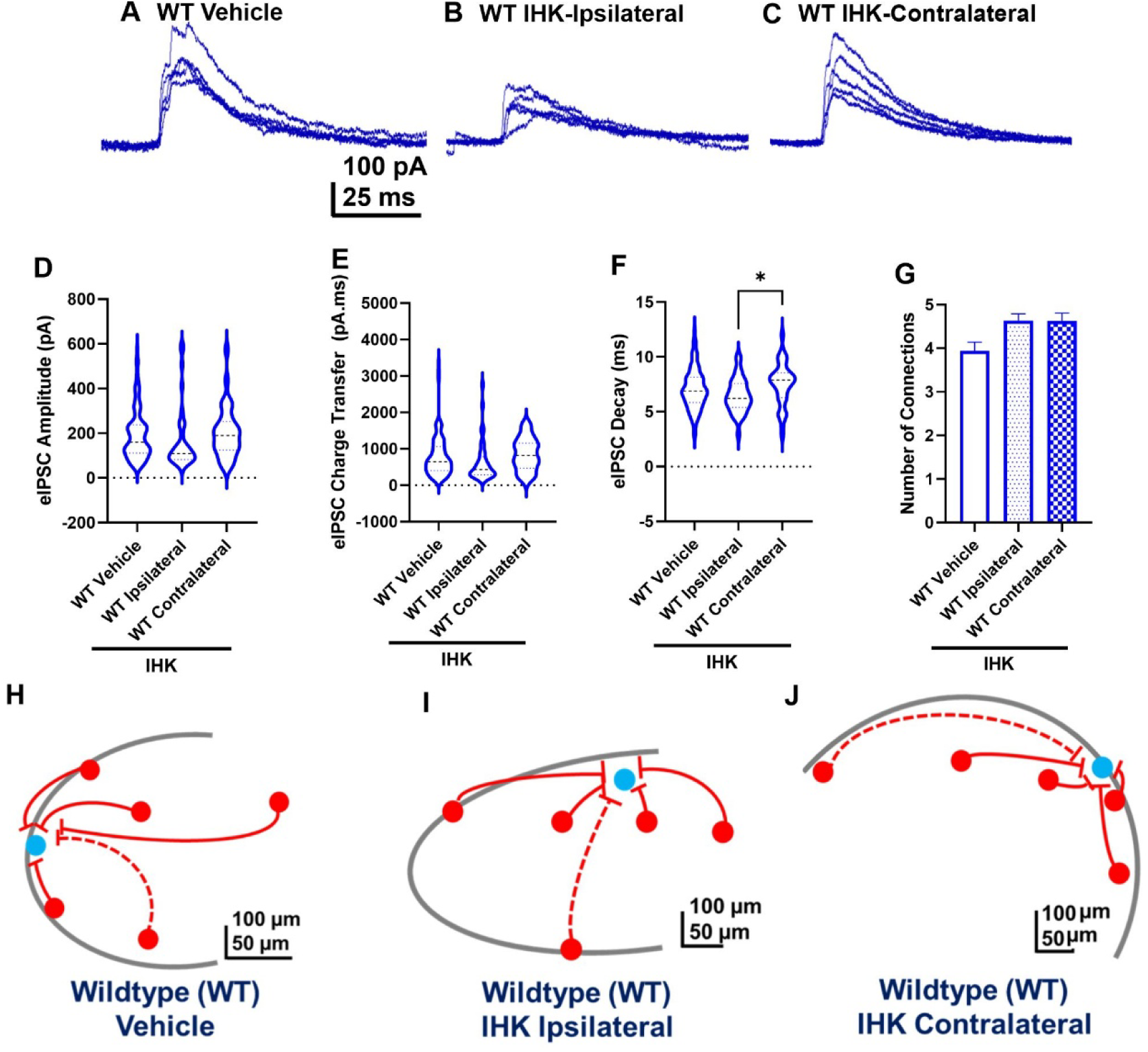
After development of TLE in wildtype mice, optogenetic stimulation of inhibitory interneurons does not reveal altered synaptic inhibition in vDGCs. (A) Representative trace showing evoked IPSCs (eIPSCs) from stimulated interneurons recorded in vDGCs from vehicle-treated wildtype (WT) mice using the stimulation protocol (5 stimulations, 10 ms duration, 1 sec interstimulus interval). The 5 eIPSCs from one stimulation train were overlayed. (B) Representative eIPSC trace from interneurons ipsilateral IHK injection in IHK-treated wildtype mice. (C) Representative eIPSC traces from interneurons contralateral to IHK injection in IHK-treated wildtype mice. Vehicle: N=10, n=123 interneurons; WT IHK: N=6, ipsilateral: n=44 interneurons, contralateral: n=26 interneurons). (D) Box and whisker plot of the peak average eIPSC amplitude across 5 stimulations from vehicle and IHK-treated wildtype mice (vehicle vs. ipsilateral: p=0.4259, vehicle vs. contralateral: p=0.9131, ipsilateral vs. contralateral: p=0.3764; One-way ANOVA). (E) Box and whisker plot of the average eIPSC charge transfer across 5 stimulations from vehicle and IHK-treated wildtype mice (vehicle vs. ipsilateral: p=0.2847, vehicle vs. contralateral: p=0.8507, ipsilateral vs. contralateral: p=0.2209; One-way ANOVA). (F) Box and whisker plot of the average decay of eIPSCs across 5 stimulations from vehicle and IHK-treated wildtype mice (vehicle vs. ipsilateral: p=0.1927, vehicle vs. contralateral: p=0.3537, ipsilateral vs. contralateral: p=0.0.323; One-way ANOVA). (G) Number of successful interneuron-vDGC connections observed from vehicle and IHK-treated wildtype mice (p>0.05, One-way ANOVA). Vehicle: N=10, n=33 vDGCs; WT IHK: N=6, ipsilateral: n=11 vDGCs, contralateral: n=8 vDGCs) The max number of interneuron connections possible for one vDGCs was 5. (H) Representative schematic of connections between 5 interneurons and a vDGC from a wildtype vehicle slice, (I) IHK-treated ipsilateral slice, and (J) IHK-treated contralateral slice. Solid lines represent successful connections, and dashed lines represent no connection.

Results suggest an increase in inhibitory synaptic input to vDGCs after development of TLE in the absence of tau expression in mice (Fig. 4; Table 3). We hypothesized that the increase in inhibitory tone involved increased connectivity of interneurons with vDGCs. Using the same optogenetic stimulation paradigm, results revealed significantly greater average peak eIPSC amplitude from interneuron connections with vDGCs ipsilateral and contralateral to IHK injection in tau^-/-^ mice that developed TLE compared to vehicle controls (Fig. 7A-D; p<0.05). Further analysis revealed numerical increases in the average charge transfer (p=0.1945) and decay of eIPSCs (p=0.0713) ipsilateral to IHK injection compared to vehicle controls, while there was a significant increase in charge transfer and decay time of eIPSCs contralateral to the IHK injection compared to vehicle controls (Fig 7E-F; p<0.05). Interestingly there were significantly more successful synaptic connections between interneurons and vDGCs ipsilateral and contralateral to IHK injection in tau^-/-^ mice compared to vehicle controls (Fig. 7G-J; p<0.05). Together, this suggests that the magnitude and number of synaptic connections between interneurons and vDGCs is greater after the development of TLE in mice lacking tau expression, which could be a result of an increase in the number of interneurons forming synapses onto a single vDGC.

**Figure 7.**
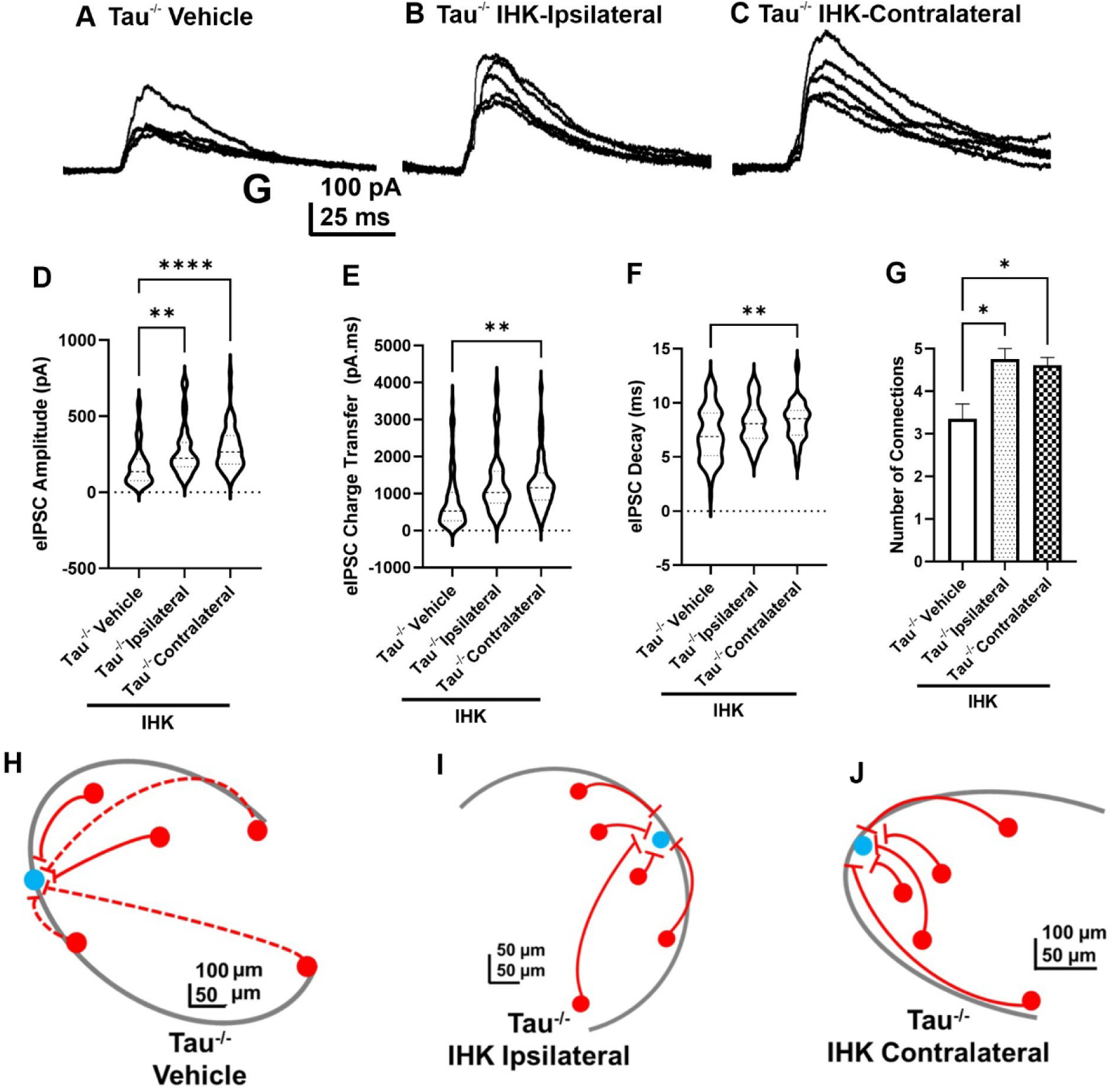
After development of TLE in tau^-/-^ mice, optogenetic stimulation of inhibitory interneurons results in greater evoked inhibition of vDGCs. (A) Representative trace showing evoked IPSCs (eIPSCs) from stimulated interneurons recorded in vDGCs from vehicle-treated tau^-/-^ mice using the stimulation protocol (5 stimulations, 10 ms duration, 1 sec interstimulus interval). The 5 eIPSCs from one stimulation train were overlayed. (B) Representative eIPSC trace from interneurons ipsilateral IHK injection in IHK-treated tau^-/-^ mice. (C) Representative eIPSC traces from interneurons contralateral to IHK injection in IHK-treated tau^-/-^ mice. Vehicle: N=10, n=64 interneurons; Tau^-/-^ IHK: N=6, ipsilateral: n=58 interneurons, contralateral: n=59 interneurons). (D) Box and whisker plot of the peak average eIPSC amplitude across 5 stimulations from vehicle and IHK-treated tau^-/-^ (vehicle vs. ipsilateral: **p=0.0030, vehicle vs. contralateral: ****p<0.0001, ipsilateral vs. contralateral: p=0.8499; One-way ANOVA). (E) Box and whisker plot of the average eIPSC charge transfer across 5 stimulations from vehicle and IHK-treated tau^-/-^ mice (vehicle vs. ipsilateral: p=0.1945, vehicle vs. contralateral: **p=0.0010, ipsilateral vs. contralateral: p=0.4280; One-way ANOVA). (F) Box and whisker plot of the average decay of eIPSCs across 5 stimulations from vehicle and IHK-treated tau^-/-^ mice (vehicle vs. ipsilateral: p=0.0713, vehicle vs. contralateral: **p=0.0041, ipsilateral vs. contralateral: p=0.8753; One-way ANOVA). (G) Number of successful interneuron-vDGC connections observed from vehicle and IHK-treated tau^-/-^ mice (p<0.05, One-way ANOVA). Vehicle: N=10, n=20 vDGCs; Tau^-/-^ IHK: N=6, ipsilateral: n=8 vDGCs, contralateral: n=13 vDGCs). The maximum number of interneuron connections possible for one vDGC was 5. (H) Representative schematic of connections between 5 interneurons and a vDGC from a wildtype vehicle slice, (I) IHK-treated ipsilateral slice, and (J) IHK-treated contralateral slice. Solid lines represent successful connections, and dashed lines represent no connection.

### High frequency stimulation of GABAergic synapses reveals changes in short term synaptic plasticity in wildtype mice after development of TLE but not in mice lacking tau expression

Next, we examined the functionality of successful inhibitory connections, by challenging positive connections with high-frequency stimulation (10 ms duration, 10 stimulations, 10 ms interstimulus interval). Analysis of the percent change in the evoked amplitude following the first stimulation revealed a lower percent change in amplitude of eIPSCs in contralateral connections in IHK-treated wildtype mice versus vehicle treated controls. This suggests a potential change in short-term plasticity involving presynaptic mechanisms in the GABAergic synapses contralateral to IHK but not ipsilateral (Fig. 8A1-4; p<0.05). Notably, there was no difference detected in the number of evoked action potentials in interneurons associated with light stimulations between interneurons from vehicle- and IHK-treated wildtype mice, suggesting similar ChR2 expression and activity in the interneurons (Supplemental Figure 2). Together, this suggests that after development of TLE in wildtype mice, there is no obvious change in the number of interneurons connected to a particular granule cell or magnitude of GABAergic synapses at sites distal from IHK injection but sustained evoked GABA release from synapses onto vDGCs across high frequency patterns of activity resulted in decreased responses after TLE development in the contralateral hemisphere of wildtype mice.

**Figure 8.**
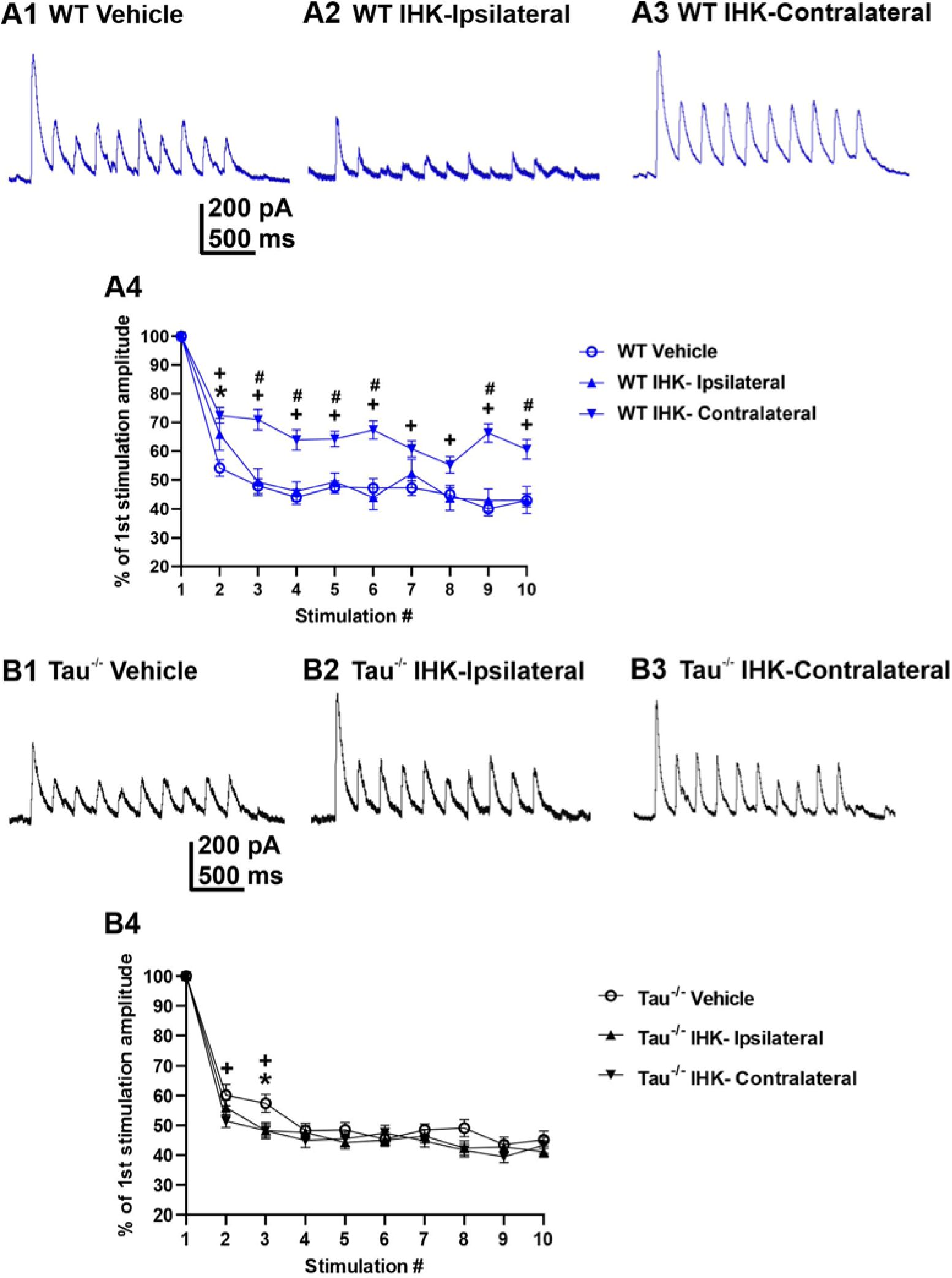
Development of TLE in wildtype mice results in sustained synaptic inhibition of vDGCs during high frequency stimulation that does not occur in tau^-/-^ mice. (A1) Representative trace showing evoked IPSCs (eIPSCs) from stimulated interneuron recorded in vDGCs from vehicle-treated wildtype (WT) mice using the stimulation protocol (10 stimulations, 10 ms duration, 10 ms interstimulus interval). (A2) Representative eIPSC trace from interneuron ipsilateral IHK injection in IHK-treated wildtype mice. (A3) Representative eIPSC trace from interneuron contralateral to IHK injection in IHK-treated wildtype mice. (A4) Summary of percent of 1^st^ pulse amplitude from connected interneuron and vDGCs challenged with high frequency stimulation. * vehicle vs. IHK ipsilateral, ^+^ vehicle vs. IHK contralateral, # IHK ipsilateral vs. IHK contralateral. Vehicle: N=9, n=27 interneurons (p<0.05, Two-way ANOVA). WT IHK: N=5, ipsilateral: n=44 interneurons, contralateral: n=34 interneurons). (B1) Representative trace showing evoked IPSCs (eIPSCs) from stimulated interneuron recorded in vDGCs from vehicle-treated tau^-/-^ mice using the stimulation protocol (10 stimulations, 10 ms duration, 10 ms interstimulus interval). (B2) Representative eIPSC trace from interneuron ipsilateral IHK injection in IHK-treated tau^-/-^ mice. (B3) Representative eIPSC trace from interneuron contralateral to IHK injection in IHK-treated wildtype mice. (B4) Summary of percent of 1^st^ pulse amplitude from connected interneuron and vDGCs challenged with high frequency stimulation. * vehicle vs. IHK ipsilateral, ^+^ vehicle vs. IHK contralateral, # IHK ipsilateral vs. IHK contralateral. Vehicle: N=9, n=50 interneurons (p<0.05, Two-way ANOVA). Tau^-/-^ IHK: N=7, ipsilateral: n=40 interneurons, contralateral: n=47 interneurons).

Differently, in tau^-/-^ mice there were no observed differences in the change in eIPSC amplitude across high frequency stimulations after TLE development. Vehicle treated mice demonstrated significantly lower percent change in amplitude in the second and third stimulations compared to ipsilateral and contralateral stimulations (Fig. 8B1-4), but unlike in wildtype mice, plasticity differences between vehicle and IHK treated tau^-/-^ mice was negligible for subsequent responses. Together these results indicate that development of TLE in mice lacking tau expression is associated with increased convergence and strengthened connectivity of inhibitory interneurons onto vDGCs distant from the focal injection of kainate that is not observed in wildtype mice that develop TLE.

## Discussion

Genetic deletion or suppression of the microtubule associated protein, tau has been shown to reduce seizure burden in models of genetic epilepsies and to suppress evoked seizures in otherwise healthy rodents (DeVos *et al*., 2013; Holth *et al*., 2013; Gheyara *et al*., 2014; Shao *et al*., 2022). The etiologies between genetic and acquired TLE models are different, in that spontaneous seizures in acquired epilepsies develop after a delay following an insult compared to seizures in genetic epilepsies that are a consequence of, for example, voltage-gated ion channel mutations. We have shown that genetic deletion of tau reduces the likelihood of evoked seizures and inhibits seizure severity but does not prevent the epileptogenic process in the IHK model of TLE (Moseley *et al*., 2025). Therefore, lack of tau expression does not prevent the epileptogenic process from occurring but is hypothesized to attenuate network activity of individual spontaneous seizures. Tau^-/-^ mice that develop TLE not only exhibit inhibitory synaptic reorganization, resulting in increased inhibitory synaptic tone in DGCs at sites distant from the initial focal insult, but also do not exhibit widespread excitatory synaptic neuroplastic changes that are associated with epileptogenesis in this and similar TLE models and in human TLE (Franck *et al*., 1995; Buckmaster *et al*., 2009; Buckmaster *et al*., 2022). Further, we uncovered previously unappreciated functions of murine tau in normal excitatory and inhibitory synaptic transmission in the dentate gyrus circuit that could support resistance to evoked seizures. Our results demonstrate a potential mechanism within the dentate gyrus local circuitry by which lack of tau expression may modify the development of TLE and reduce spontaneous seizure burden in a manner consistent with synaptic reorganization of inhibitory circuitry.

In rodent models of acquired TLE, epilepsy development promotes hyperactivity of DGCs that may result from recurrent excitatory synaptic connections between DGCs linked to sprouting of axon collaterals of DGCs themselves (Cronin *et al*., 1992; Althaus *et al*., 2015; Kelly & Beck, 2017). Few studies have examined excitability changes of DGCs associated with TLE using the IHK model. Interestingly, our results demonstrate a significant decrease in excitability of dDGCs near the IHK injection where severe hippocampal damage and DGC dispersion are detected (Moseley *et al*., 2025). It is possible that development of spontaneous seizures lowers the excitability of DGCs near the focal injury, suggesting that spontaneous seizures initiate distal to the injury site in this model. Differently, we found that development of TLE in the IHK model is associated with an increase in action potential firing frequency of DGCs in the ventral dentate gyrus, regardless of tau expression. Together these results highlight an association between TLE development and increased excitability of DGCs at sites distal to the focal injury, indicating a potential widespread effect of epileptogenesis on neuronal excitability throughout the dentate gyrus.

Development of spontaneous seizures is associated with an increase in excitatory synaptic input to DGCs in the dorsal dentate gyrus, as we and others have shown in this and other rodent models of acquired TLE (Cronin *et al*., 1992; Buckmaster & Dudek, 1997; Wuarin & Dudek, 2000; Sutula & Dudek, 2007; Hunt *et al*., 2010; Althaus *et al*., 2016; Moseley *et al*., 2025). In the context of tau deletion and epileptogenesis, the increase in excitatory input to dDGCs ipsilateral to the IHK injection was similar between the two genotypes, even though mice lacking tau expression exhibit less severe spontaneous seizure expression (Moseley *et al*., 2025). This supports a role for excitatory synaptic reorganization in the evolution of spontaneous seizure development in the IHK model, but it does not intuitively account for the difference in expression of individual seizures across the genotypes once TLE has developed. Here, we found no detectable change in excitatory input to vDGCs in wildtype or mice lacking tau expression after epileptogenesis. This suggests an attenuation of the TLE associated excitatory synaptic reorganization in the ventral dentate gyrus in tau^-/-^ mice, which could contribute to lower seizure expression in this genotype. Interestingly, there was no change in stochastic glutamatergic release (i.e., mEPSC frequency) regardless of epileptogenesis or tau expression, suggesting the observed increase in sEPSC frequency contralateral compared to ipsilateral to IHK may be due to activity of afferent excitatory neurons (e.g., entorhinal cortex or hilar mossy cells) in mice expressing endogenous tau protein, which was not observed in the absence of tau expression. Excitatory synaptic reorganization of DGCs is associated with MFS after TLE development, where sprouted axons of DGCs can synapse onto other DGCs, increasing their overall excitatory synaptic input. While MFS was detected in mice that developed TLE, we did not detect differences in MFS between genotypes. Differences in sprouted mossy fiber synapse maturity or in the dynamics of the synapses themselves may account for the mismatch between the anatomical and electrophysiological outcomes, since our results demonstrate no change in overall excitatory synaptic input to DGCs in the ventral hippocampus that occurs once TLE develops in the IHK model. Further, DGCs in the ventral dentate gyrus in naïve tau^-/-^ mice receive less overall excitatory synaptic input compared to wildtype controls, highlighting a possible role of tau protein in glutamatergic signaling upstream of the dentate gyrus through activity of afferent excitatory neurons (e.g., entorhinal cortex or hilar mossy cells) that may be associated with lower susceptibility to evoked seizure expression and SE that we and others have observed in tau^-/-^ mice (Li *et al*., 2014; Putra *et al*., 2020; Shao *et al*., 2022; Moseley *et al*., 2025).

Seizures in rodent models of TLE often result in a significant loss of inhibitory synaptic input to DGCs that is associated with a loss of select groups of inhibitory interneurons (Buckmaster & Dudek, 1997; Kobayashi & Buckmaster, 2003; Hunt *et al*., 2011; Butler, 2017; Moseley *et al*., 2025). We recently showed that development of TLE is associated with an increase in inhibitory synaptic input in dDGCs contralateral to the IHK injection in mice lacking tau expression, suggesting that inhibitory synaptic reorganization is possible at sites outside the focal injury (Moseley *et al*., 2025). It has been proposed that increased synaptic excitation of GABAergic interneurons due to sprouted mossy fibers or other afferents synapsing on interneurons in the dentate gyrus may partially compensate for interneuron loss in rodent models of acquired TLE and post-traumatic epilepsy (Sloviter, 1992; Hunt *et al*., 2011; Butler *et al*., 2015; Kang *et al*., 2022). In this study, we found that overall inhibitory input was preserved in the ventral dentate gyrus in wildtype mice, suggesting survival of inhibitory interneurons distal to the IHK injection, even in the ipsilateral hemisphere. Previous studies demonstrated a reduction in inhibitory synaptic input to vDGCs and pyramidal cells in the CA1 after development of TLE in the IHK model (Kang *et al*., 2021; Lee *et al*., 2024). Differently, while overall inhibitory synaptic input was unchanged in wildtype mice here, we found a decrease action-potential independent inhibitory synaptic events after development of TLE, consistent with a loss of functional inhibitory synapses onto vDGCs in wildtype mice. Conversely, an increase in inhibitory synaptic input to vDGCs was detected after development of TLE in mice lacking tau expression, both ipsilateral and contralateral to the IHK injection. Together with findings from the dorsal dentate gyrus, this suggests that epileptogenesis results in increased inhibitory synaptic signaling at sites distant from the focal injury in tau^-/-^ mice that does not occur in wildtype mice (Moseley *et al*., 2025). The increase in synaptic inhibition, combined with the relatively lowered emergent synaptic excitation in vDGCs may contribute to the lower spontaneous seizure expression in tau^-/-^ mice, even after TLE development.

Increased inhibitory synaptic input to DGCs has been associated with axon sprouting of surviving inhibitory interneurons (Halabisky *et al*., 2010) and could also involve increased DGC input to surviving interneurons (Kang *et al*., 2021; Kang *et al*., 2022), thereby increasing their overall output onto DGCs (Sloviter, 1992; Kotti *et al*., 1997; Wenzel *et al*., 2000; Buckmaster *et al*., 2002; Cavazos *et al*., 2003; Sloviter *et al*., 2006). We used an AAV construct with ChR2-mCherry fusion that drives selective expression in GABAergic interneurons under the control of the Dlx promoter (Dimidschstein *et al*., 2016). To date, there are no studies indicating a differential role of tau protein among specific interneuron subtypes (Shao *et al*., 2022). Injection of the AAV in the ventral dentate gyrus, resulted in robust expression in all mice (wildtype and tau^-/-^) with no obvious expression differences detected between vehicle-treated and IHK-treated mice. Since we detected no differences in sEPSC frequency or amplitude across treatments or genotypes in mCherry-labeled inhibitory interneurons, we focused on connections between interneurons and vDGCs to assess the differences in synaptic inhibition of vDGCs across genotypes after TLE development.

Decreased inhibitory synaptic input suggests a lower probability of GABA release and/or reduced number of functional GABAergic synapses contacting vDGC neurons in wildtype mice after TLE development. However, we did not observe significant differences in eIPSC amplitude, charge transfer, decay time constant, or the number successful connections between activated interneurons and vDGCs after TLE development in wildtype mice compared to vehicle controls. Increasing the number of interneurons stimulated, either separately or concurrently, might reveal potential subtle connectivity changes with TLE in wildtype mice, but significant changes in inhibitory synaptic connectivity associated with TLE development were not detected in vDGCs from wildtype mice. Nevertheless, optogenetic stimulation of inhibitory interneurons in tau^-/-^ mice resulted in significantly increased average eIPSC amplitude, charge transfer, and decay time constant, which coincided with an increased number of successful connections detected between interneurons and recorded vDGCs after TLE development. Reactive synaptogenesis of surviving interneurons has been proposed as a compensatory mechanism that restores at least some part of synaptic inhibition that may be lost during TLE development. In resected human tissue and in rodent models of TLE, GABAergic terminals of somatostatin-expressing interneurons express immunoreactivity throughout the molecular layer of the dentate gyrus that is initially decreased shortly following status epilepticus followed by a persistent increase during and after epileptogenesis (Davenport *et al*., 1990; deLanerolle *et al*., 1992a; deLanerolle *et al*., 1992b; Mathern *et al*., 1995; Houser & Esclapez, 1996; Buckmaster & Dudek, 1997; Mathern *et al*., 1997; Andre *et al*., 2001; Kobayashi & Buckmaster, 2003). In tissue from patients with TLE, other interneurons that provide perisomatic inhibition to DGCs increase axon basket formation surrounding DGCs and their axon initial segments (Arellano *et al*., 2004). Moreover, cholecystokinin expressing interneurons not only increase synapses onto axon DGC initial segments of but also synapse in the molecular layer of the dentate gyrus after TLE develops (Gruber *et al*., 1993; Wittner *et al*., 2001). It is possible that the increased inhibitory synapse formation into DGCs after TLE development observed here contributes to the lower spontaneous seizure burden in tau^-/-^ mice. These results also indicate a potential role for tau protein in inhibitory synaptogenesis normally, since synaptic inhibition was lower in vehicle treated tau^-/-^ than in wildtype mice.

In addition to modified inhibitory input to DGCs and reactive inhibitory synaptogenesis, modulation of inhibitory synaptic plasticity is also associated with TLE in the dentate gyrus. Here, we measured short-term plasticity of GABAergic synapses using high frequency optogenetic stimulation of inhibitory interneurons, finding evidence of potentially compensatory synaptic plasticity of surviving inhibitory inputs to vDGCs in wildtype, but not tau^-/-^ mice after TLE development. These findings suggest that short-term plasticity of GABAergic synapses occurs in association with TLE development in wildtype mice, which is not present in mice lacking tau expression. Combined with the observed increase in eIPSC responses and increase in interneuron connectivity, it is likely that development of TLE in mice lacking tau expression is associated with significant synaptic reorganization of inhibitory interneurons at sites distant to the initial focal kainate injection that does not occur in wildtype mice.

## Conclusions

In this study we have shown that neuroplastic changes associated with TLE development after dorsal IHK injection are detectable in the ventral dentate gyrus. Additionally, we identified potential synaptic mechanisms involving distant inhibitory synaptic reorganization that may impact the degree of protection from seizures but not from the process of epileptogenesis in tau^-/-^ mice. Being that tau protein can regulate synapses through microtubule stability, protein translation, and activity-dependent remodeling (Chang *et al*., 2021; Thomas *et al*., 2025), it is possible that when tau is not present to act as a “brake” on synaptic reorganization the homeostatic constraints that exist in situations of increased excitability allow new inhibitory connections to form more readily. The finding that interneurons and synaptic inhibition appear to be overtly affected by tau deletion is at odds with previous work highlighting that knockout of tau in excitatory neurons globally, and not inhibitory neurons, reduced seizure activity (Shao *et al*., 2022). It is possible that DGCs are not affected by the loss of tau as other excitatory neurons are since DGCs are unique in their slow firing during activation compared to other excitatory cell types in the hippocampus. With the output regulation of the dentate gyrus being tightly controlled by synaptic inhibition, it is conceivable that a lack of tau expression impacts overall dentate gyrus circuit activity primarily through the inhibitory pathway in the dentate gyrus. Further studies are required to elucidate modifications of specific receptor mediated synaptic plasticity and the contribution of specific inhibitory neuron subtypes to the changes in synaptic inhibition in tau^-/-^ mice and whether inhibitory synaptic reorganization reflects a compensatory outcome or is an emergent quality of TLE epileptogenesis.

## Acknowledgements

This work was funded by NIH NINDS R01 NS092552

## Author Contributions

M.C.M and B.N.S. designed research; M.C.M performed research; M.C.M. analyzed data; M.C.M. and B.N.S wrote the paper

---

**Supplemental Figure 1:**
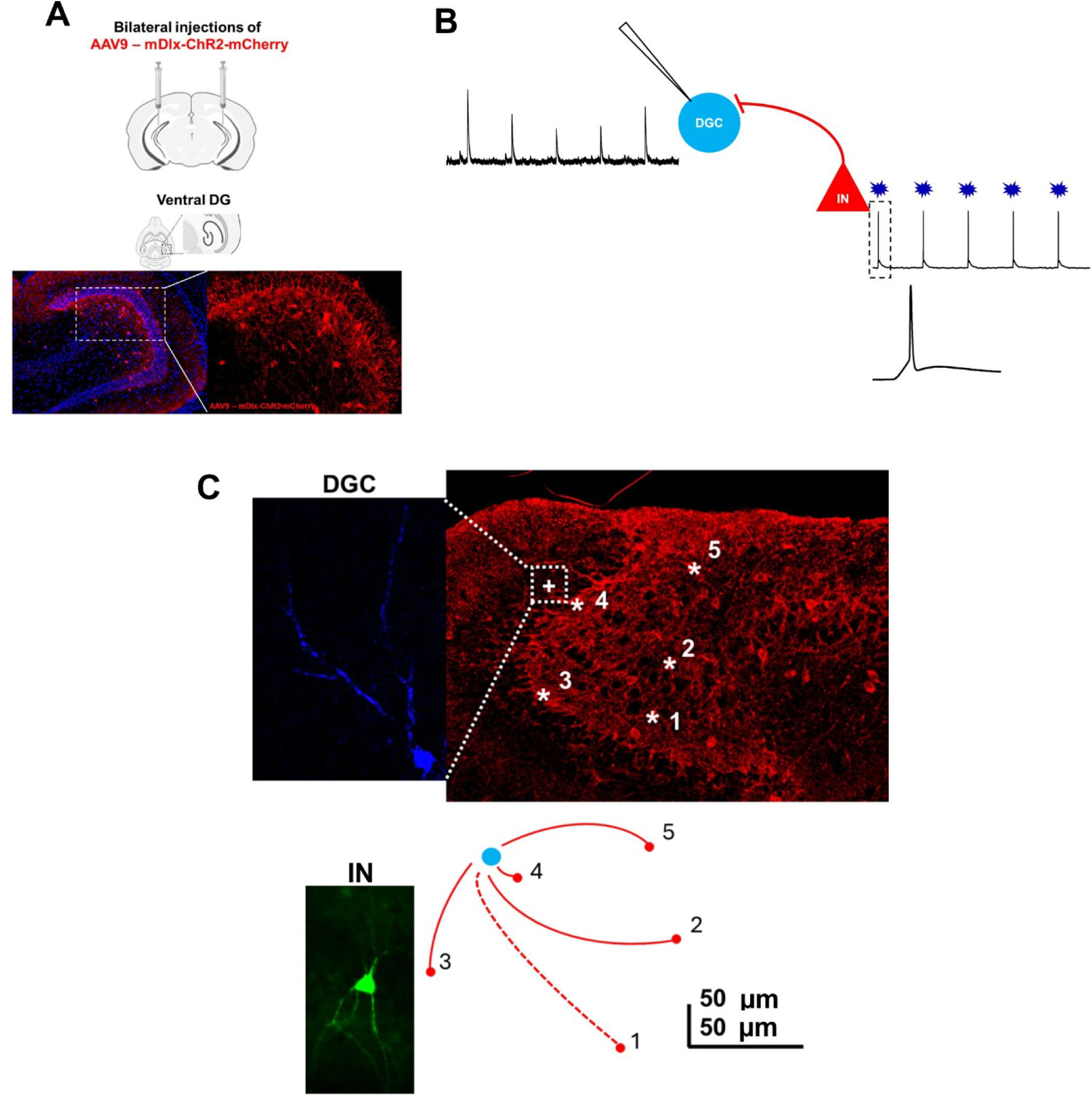
Experimental design of optogenetic stimulation of ChR2-mCherry expressing interneurons. (A) Bilateral injections of AAV9-mDlx-ChR2-mCherry virus into the ventral dentate gyrus (DG) results. Representative image of viral expression (red) and DAPI (blue). (B) Optogenetic stimulation of interneurons and recording DGCs. 5 blue light stimulations (10 ms duration, 1 sec intervals) elicits action potentials in ChR2 expressing interneurons. Successful connections between stimulated interneuron and DGC results in evoked IPSCs corresponding to action potentials. (D) Experimental design. Biocytin filled DGC, recorded at + positive on slice. Representative image of ventral DG slice with identified stimulated interneurons (*). Schematic of actual micron distance of recorded DGC (blue) and stimulated interneurons (red: 1-5). Solid red line represents successful connections (2-5). Dashed red line represents unsuccessful connection (1). Image of biocytin filled (3) interneuron.

**Supplemental Figure 2:**
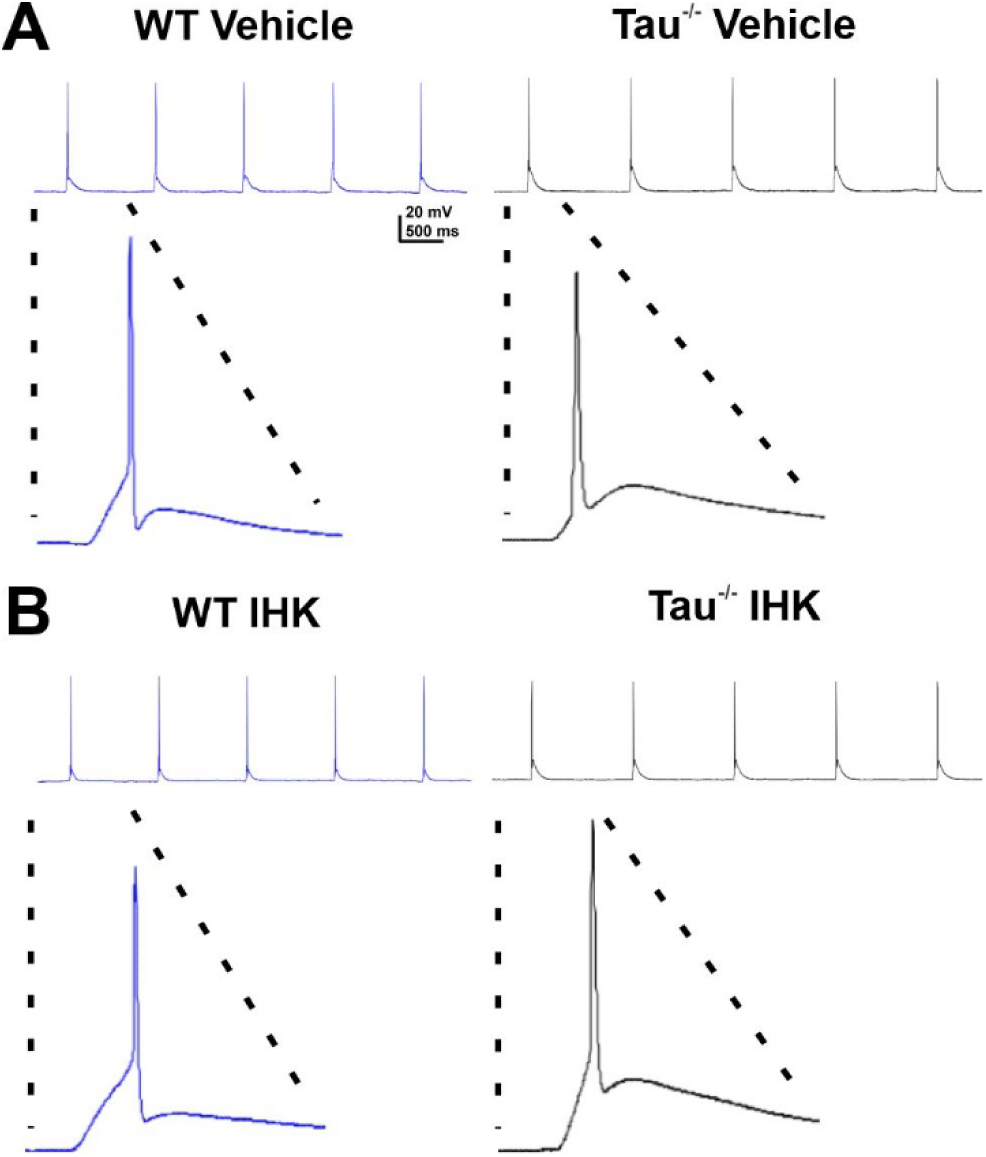
Optogenetic stimulation reliably evoked action potentials in ChR2-expressing interneurons. (A) Representative traces of light-evoked action potentials (APs) in ChR2-expressing interneurons from vehicle treated WT and tau^-/-^ mice. (B) Representative traces of light-evoked action potentials in ChR2-expressing interneurons from IHK-treated WT and tau^-/-^ mice. The first AP from each trace is shown at a higher resolution. Each blue light stimulation reliably evoked one AP in each interneuron.

